# Virus-like particles enable targeted gene engineering and pooled CRISPR screening in primary human myeloid cells

**DOI:** 10.64898/2025.12.14.692434

**Authors:** Hyuncheol Jung, Pascal Devant, Carter Ching, Mineto Ota, Jennifer Hamilton, Zachary Steinhart, Wayne Ngo, Luis Sandoval, Jae Hyung Jung, Da Xu, Meirui An, Esha Urs, Peixin Amy Chen, Vincent Allain, Takuya Tada, James K Nuñez, Nathaniel R Landau, David R. Liu, Justin Eyquem, Jennifer A. Doudna, Alexander Marson, Julia Carnevale

**Affiliations:** Gladstone-UCSF Institute for Genomic Immunology, San Francisco, CA 94158, USA; Department of Medicine, University of California San Francisco, San Francisco, CA 94143, USA; Parker Institute for Cancer Immunotherapy, San Francisco, CA 94129, USA; UCSF Helen Diller Family Comprehensive Cancer Center, University of California, San Francisco, San Francisco, CA 94158, USA; Institute for Human Genetics (IHG), University of California, San Francisco, San Francisco, CA 94143, USA; Department of Microbiology and Immunology, University of California, San Francisco, San Francisco, CA 94143, USA; Chan Zuckerberg Biohub San Francisco, San Francisco, CA, USA; Gladstone Institute for Data Science and Biotechnology; San Francisco, CA 94158, USA; Innovative Genomics Institute, University of California, Berkeley, Berkeley, CA, USA; Department of Molecular and Cell Biology, University of California, Berkeley, Berkeley, CA, USA; California Institute for Quantitative Biosciences, University of California, Berkeley, Berkeley, CA, USA; Howard Hughes Medical Institute, University of California, Berkeley, Berkeley, USA; Molecular Biophysics and Integrated Bioimaging Division, Lawrence Berkeley National Laboratory, Berkeley, CA, USA; Department of Chemistry, University of California, Berkeley, Berkeley, CA, USA; Department of Microbiology, NYU Grossman School of Medicine, New York, NY, 10016, USA; Department of Genetics, Stanford University, Stanford, CA; Merkin Institute of Transformative Technologies in Healthcare, Broad Institute of MIT and Harvard, Cambridge, MA, USA; Department of Chemistry and Chemical Biology, Harvard University, Cambridge, MA, USA; Howard Hughes Medical Institute, Harvard University, Cambridge, MA, USA; Université Paris Cité, INSERM UMR1342, Hôpital Saint-Louis, Paris, France

**Author notes:** These authors contributed equally (Hyuncheol Jung, Pascal Devant). Present address: Azalea Therapeutics (Jennifer Hamilton).

## Abstract

Primary human myeloid cells are promising candidates for immunotherapy, yet efficient and scalable technologies for genetic engineering and screening in these cells are limited. Here we present a virus-like particle (VLP)-based toolkit that delivers diverse CRISPR genome editing modalities to human monocytes, macrophages, and dendritic cells with high efficiency while preserving viability and innate immune responsiveness. VLP-mediated delivery of ribonucleoprotein payloads supports gene knockout, base editing and epigenetic silencing, and enables site-specific integration of large DNA sequences when combined with AAV donors for homology-directed repair. Leveraging sgRNA delivery via VPX-lentivirus combined with Cas9 protein delivery via engineered virus-like particle (eVLP) treatment (“SLICeVLP”), we performed the first pooled loss-of-function screens in human macrophages. We uncovered regulators of TNF production and CD80 expression in human macrophages, converging on TNFAIP3 as a central regulator of inflammatory polarization. TNFAIP3 ablation promoted a pro-inflammatory cell state that is resistant to suppressive polarization, and augmented cytotoxicity of engineered HER2 CAR-macrophages. Taken together, this technology platform enables unbiased discovery and characterization of functional gene targets in primary human myeloid cells.

## INTRODUCTION

The genetic manipulation of primary human myeloid cells has emerged as a promising avenue for both basic research and therapeutic intervention in immune-mediated diseases. Myeloid populations, including monocytes, macrophages, and dendritic cells, play critical roles in innate immunity, antigen presentation, and the orchestration of immune responses. These cells readily home to sites of infection, inflammation, or tumors and have evolved to sense and modulate the local microenvironment. These features make myeloid cells attractive targets for immunotherapies aimed at enhancing pathogen clearance, eliminating malignant cells or regulating inflammation^1–3^.

There is a critical need for genetic screening platforms to discover regulators of myeloid cell polarization, activation, and survival. Such platforms would facilitate the systematic identification of novel gene targets, thereby improving efficacy and safety of myeloid cell-based therapies. To date, all functional myeloid cell CRISPR screening studies have been performed in immortalized human cell lines or primary murine cells^4–11^, which incompletely recapitulate human myeloid cell biology, or have employed arrayed screening technologies^12^, which are inherently limited in scalability and can be challenging to implement without specialized equipment for laboratory automation. The lack of scalable methodologies for pooled high-throughput screening in primary human myeloid cells has left numerous biological mechanisms and potential therapeutic targets unexplored, thereby slowing progress toward advanced myeloid-based cell therapies. In this context, the efficient delivery of genetic engineering payloads, such as different Cas9 variant genes or proteins, into primary human myeloid cells represents a major hurdle. Therefore, there is a pressing need for efficient delivery systems that maintain cell viability and functionality, facilitating robust functional genomics analyses in primary human myeloid cells.

Recent pre-clinical studies demonstrated that macrophages engineered to express chimeric antigen receptors (CARs) can invade tumors, phagocytose cancer cells and activate adaptive immune responses through cross-presentation and epitope spreading^13–15^. These findings underscore the potential of myeloid-directed therapies to overcome the immunosuppressive tumor microenvironment (TME) and the challenges posed by solid tumors^16^. A first clinical trial leveraging CAR-macrophages similarly reported successful infiltration of engineered macrophages into tumor sites without dose-limiting toxicities, severe cytokine release syndrome, or immune effector cell-associated neurotoxicity syndrome^17^. Despite this progress, the impact of engineered myeloid cells on patient survival and tumor control has remained modest to date, highlighting a need for new strategies to improve these therapies. Myeloid cells are highly plastic, and within the TME they are readily polarized into cell states that support cancer progression and suppress the adaptive immune system^18^. Developing strategies to steer their polarization states in situ away from suppressive and into immunostimulatory states will likely greatly enhance myeloid cell therapies^16^. One promising approach to enhance the efficacy of adoptive cell therapies is through engineering of cell functions by means of genome editing^19^. In addition to engineering homing and targeting of myeloid cells with CARs or other synthetic receptors, great potential could be unlocked if we can simultaneously engineer signaling pathways and cellular responses through genome or epigenome reprogramming. This would require an expanded set of tools for genetic and/or epigenetic programming of myeloid cell functions. CRISPR-based tools for targeted gene knock-outs, base edits and epigenetic silencing are now being increasingly used as strategies to enhance T cell therapies^20–24^, but have not yet been extended broadly to myeloid cell engineering, thereby limiting our ability to fully exploit their therapeutic potential.

Here, we present a toolbox for the genetic engineering of human primary myeloid cells, which relies on the delivery of CRISPR gene editing enzymes and sgRNAs through virus-like particle (VLP)-based approaches. We demonstrate that VLP-based strategies can be used for high-efficiency gene knockout, site-specific knock-in, base editing, as well as epigenetic gene silencing, while maintaining cell viability and responsiveness to common innate immune stimuli, which we show are both compromised by electroporation. Furthermore, we developed a screening platform combining lentiviral delivery of pooled sgRNA libraries with Cas9-loaded VLP treatment to enable high-throughput pooled loss-of-function screening in human macrophages. We applied this technology to identify regulators of inflammatory cytokine production and macrophage polarization, which converged on the gene *TNFAIP3* as a conserved master regulator of macrophage responses. Finally, we demonstrate that *TNFAIP3* ablation confers resistance to polarization toward immunosuppressive cell states and enhances CAR-macrophage-mediated cancer cell killing in vitro, highlighting the potential of this gene discovery pipeline to drive advancements in cell editing for therapeutic applications.

## RESULTS

### Virus-like particles achieve efficient gene editing in human myeloid cells while preserving viability and function

Introducing CRISPR editing molecules into myeloid cells has remained challenging, despite impressive progress in other primary human immune cell types. Nucleofection with Cas9 ribonucleoproteins (RNPs) can yield high gene editing efficiency in primary human myeloid cells^12,25,26^, and we were able to reproduce these findings (Extended Data Fig. 1a). However, we observed substantial cell loss and severe functional impairment among surviving cells, characterized by markedly reduced inflammatory cytokine production in response to canonical stimuli, bacterial lipopolysaccharide (LPS) and interferon-γ (IFN-γ; Extended Data Fig. 1b,c). These limitations greatly hinder the applicability of nucleofection for studying myeloid cell biology and its potential therapeutic use in CRISPR-based reprogramming of myeloid cells. RNP-loaded virus-like particle (VLP) platforms have recently emerged as promising tools for the direct delivery of gene editing enzymes into cells. One such technology, termed Cas9 Enveloped Delivery Vehicles (Cas9EDVs), involves Cas9 fused to the HIV gag protein via a protease-cleavable linker, packaged together with sgRNA into an enveloped VLP^27,28^. To assess the applicability of Cas9EDVs for genome editing in primary human myeloid cells, we generated Cas9EDVs carrying a ribonucleoprotein (RNP) complex targeting the *B2M* locus. Cas9EDV treatment of freshly isolated monocytes or differentiated macrophages resulted in efficient B2M disruption, comparable to that achieved by nucleofection. Notably, unlike nucleofection, Cas9EDV treatment largely preserved cell numbers following editing (Fig. 1a, b). More importantly, whereas nucleofected cells demonstrated clear dysfunction, EDV-edited cells retained normal functionality, producing cytokine responses to LPS/IFN-γ stimulation at levels comparable to unedited controls (Fig. 1c). As an alternative approach, we tested a Moloney Murine Leukemia Virus (MMLV)-based engineered virus-like particle (eVLP) system, conceptually similar to Cas9EDV, but noted for utilizing a less mature viral capsid of spherical shape, allowing for the packaging of a particularly high amount of RNPs into each VLP^29,30^. We used the latest generation (v5) of eVLP particles, which have been optimized by directed evolution for enhanced RNP delivery ^29,30^. This system demonstrated very high efficiency editing, leading to almost complete gene and protein disruption at higher doses tested, with minimal toxicity in monocytes and macrophages (Fig. 1d, e and Extended Data Fig. 1d,e). Similar to macrophages, we also observed dysfunction of dendritic cells after nucleofection of Cas9-RNPs (Extended Data Fig. 1f,g). In contrast, we found that the EDV-delivery approach proved highly effective in immature and mature dendritic cells, with preservation of canonical cell functions, highlighting its utility across multiple myeloid cell lineages. (Fig. 1f).

**Figure 1:**
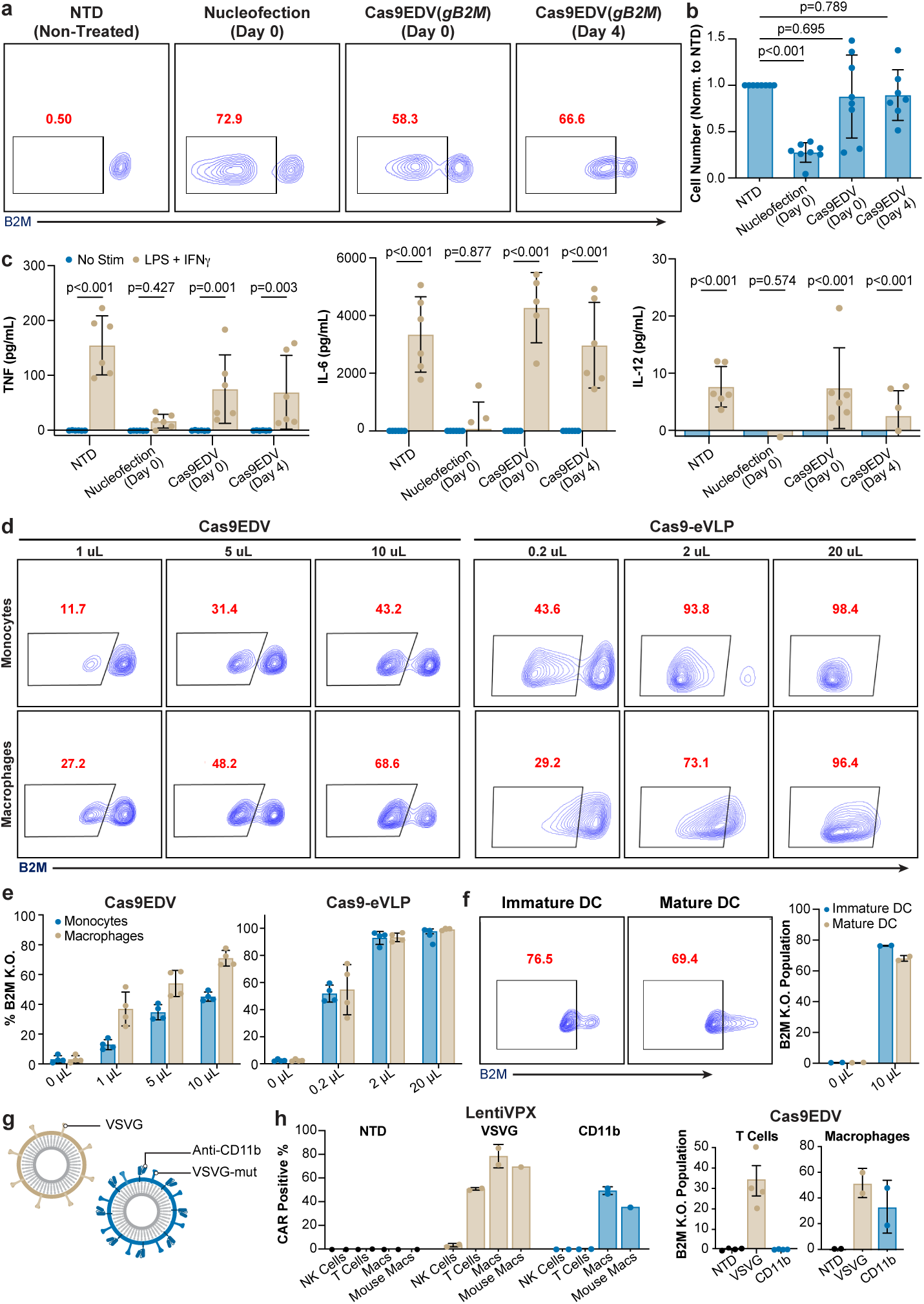
VLP-mediated delivery of CRISPR ribonucleoproteins into human primary myeloid cells enables high efficiency gene editing while preserving cell viability. **a**, Representative flow cytometry plots showing knockout of B2M protein in primary human macrophages after delivery of Cas9-RNP by nucleofection or Cas9EDV at Day 0 or Day 4 post isolation. **b**, Normalized cell yield after nucleofection or Cas9EDV treatment of primary human macrophages. *P* values calculated by means of Dunnett’s multiple comparison test after ordinary one-way ANOVA tests. **c**, Nucleofected or Cas9EDV-treated primary human macrophages were stimulated with 100 ng/ml of LPS and 25 ng/ml of IFN-γ and cytokine release was measured after 24h. *P* values calculated by means of Tukey’s multiple comparison test after ordinary two-way ANOVA tests. **d**, Representative flow cytometry plots showing knockout of B2M protein after treatment of monocytes (Day 0 post-isolation) or macrophages (after 4 days of differentiation with M-CSF) with indicated doses of Cas9EDVs or Cas9-eVLPs carrying an sgRNA targeting *B2M.* **e**, Quantification of B2M-deficient fraction of cells after Cas9EDV or Cas9-eVLP treatment of monocytes or macrophages. **f**, Representative flow cytometry plots and quantification of B2M protein knockout after treatment of immature or mature human primary dendritic cells with indicated doses of Cas9EDVs or Cas9-eVLPs. **g**, Schematic representation of CD11b-targeted Cas9EDV particles. **h**, Human primary macrophages, T cells, NK cells or murine primary macrophages were treated with VPX lentiviral vectors encoding HER2-CAR + GFP or Cas9EDVs loaded with a sgRNA targeting B2M. Particles were pseudotyped either with conventional VSVG or VSVGmut and membrane-bound scFv against CD11b. CD11b-pseudotyped particles facilitated targeted delivery of payloads into human and murine macrophages. Each data point represents the result of one independent assay. Bars and error bars represent mean ± standard deviation. Flow plots are representative of at least three independent experiments.

### Bulk RNA-seq analysis uncovers transcriptional changes following nucleofection and Cas9EDV treatment

We next examined the transcriptional differences in primary human monocytes and macrophages edited via nucleofection vs. Cas9EDV. For bulk RNA-seq analysis, peripheral blood-derived CD14⁺ monocytes were harvested one day after engineering, while macrophages were differentiated for seven days following genetic manipulation (Extended Data Fig. 2a). We targeted the *AAVS1* locus, a safe-harbor site to ensure integration with minimal impact on intrinsic cellular signaling and function (Extended Data Fig. 2b). We then performed gene set enrichment analysis (GSEA) to identify pathways differentially activated in nucleofected or Cas9EDV-treated cells compared to non-treated controls. Both engineering modalities induced an inflammatory gene signature even in the absence of specific stimulation, including upregulation of genes associated with interferon alpha, and interferon gamma signaling pathways, suggesting that some degree of stimulation may be a general feature of Cas9 or sgRNA delivery into myeloid cells (Extended Data Fig. 2c). However, nucleofected cells, unlike those treated with Cas9EDVs, exhibited specific enrichment of mTORC1 signaling, xenobiotic metabolism, and oxidative phosphorylation pathways, suggesting that nucleofection may alter the metabolic programming of myeloid cells.

To further assess the transcriptionally evident functional changes resulting from these genetic engineering strategies in myeloid cells, we conducted a second bulk RNA-seq experiment in which edited macrophages were stimulated with LPS and IFN-γ. Compared with unedited controls or Cas9EDV-treated macrophages, nucleofected macrophages demonstrated attenuated transcriptional responses to stimulation, including reduced activation of inflammatory response, interferon-gamma, and interferon-alpha signaling pathways (Extended Data Fig. 2d-f). These findings are consistent with the diminished inflammatory cytokine production observed earlier.

Because nucleofected macrophages exhibited hyporesponsiveness upon stimuli, we next examined the relationship between nucleofected macrophages and previously identified transcriptional signatures of specific macrophage subpopulations^31^. Nucleofected macrophages closely aligned with the CD14CTX program described in immune checkpoint-resistant bladder cancer, and resembled SPP1⁺ tumor-associated macrophages, a population associated with stromal remodeling and immune suppression^31–33^. Both programs are characterized by elevated expression of immunosuppressive cytokines and chemokines. In contrast, Cas9EDV treated cells most closely resembled ISG15⁺ macrophages, which are consistent with higher expression of canonical M1-like signatures^34^. Furthermore, nucleofected cells exhibited gene expression patterns consistent with a less differentiated, immature monocyte-like state, whereas Cas9EDV-treated cells displayed signatures associated with more mature, differentiated macrophages (Extended Data Fig. 2g). Taken together, these data suggest that nucleofection, but not VLP-treatment, of myeloid cells may induce an immunosuppressive phenotype and reduce their ability to respond to common inflammatory stimuli.

### Modified viral envelopes for targeting CD11b positive immune cells

VLPs not only allow for highly efficient and non-toxic delivery of Cas9-RNPs, but they also have the inherent advantage that they can be engineered to have selectivity for specific cell types. This feature was showcased previously by the specific targeting of Cas9EDVs to T cells^28^. In the experiments described above, we pseudotyped VLPs with vesicular stomatitis virus glycoprotein (VSVG), which allows for highly efficient, yet non-specific viral entry into virtually all mammalian cells, since it makes use of the ubiquitously expressed and highly conserved low-density lipoprotein receptor family as receptors^35^. We wondered whether VLPs could be designed to specifically target myeloid cells. To this end, we leveraged a mutant form of VSVG (VSVGmut), which maintains endosomal fusion activity but lacks affinity for its native target^36^. We co-expressed this mutant envelope with a membrane-bound scFv against CD11b, a marker broadly expressed on myeloid cells, during VLP packaging in HEK293T cells (Fig. 1g). Using this CD11b-scFv-pseudotyping approach, we achieved efficient and selective delivery of payloads by both lentiviral vectors and Cas9EDVs to primary human macrophages while sparing other immune cell types like T cells. Notably, since the scFv we used can bind both human and murine CD11b, we were able to specifically transduce not only human but also murine macrophages (Fig. 1h). This targeting specificity advantage highlights another potentially beneficial feature of harnessing Cas9EDVs for therapeutic engineering in myeloid cells.

### Site-specific Knock-in by combining VLPs with AAV6-mediated delivery of HDR donor templates

Recent advances in genome editing have enabled precise transgene integration into targeted genomic sites in human lymphocytes, with homology directed repair (HDR)-mediated insertion of large genetic elements transforming the design of adoptive cell therapies and supporting the production of numerous clinical products^37–39^. The use of adeno-associated virus (AAV) as a DNA donor template is one effective strategy for high-efficiency HDR knock-in efficiencies in T cells^40,41^. In contrast, HDR in primary human monocytes and macrophages remains challenging due to their predominantly post-mitotic state, innate immune activation in response to nucleic acid delivery, and the absence of robust Cas9 delivery methods. Building on our findings that Cas9EDVs achieve high gene editing efficiency in primary human macrophages, we investigated whether the combination of AAV and Cas9EDV could facilitate efficient and site-specific integration of large DNA sequences.

We first designed a strategy in which co-delivery of AAV serotype 6 (AAV6) and Cas9EDV mediates the targeted insertion of a promoter-less GFP cassette into the first exon of the clathrin light chain A (CLTA) gene by HDR (Fig. 2a). To test this approach, cells from two commonly used human myeloid cell lines (U937 and THP-1) were co-treated with AAV6 expressing a GFP-CLTA HDR template, and Cas9EDVs loaded with an sgRNA targeting a site near the start codon of in the CLTA gene, followed by analysis of GFP expression by flow cytometry. Notably, we observed GFP-positive populations of >40% in U937 cells and >80% in THP-1 cells, demonstrating robust knock-in of GFP (Fig. 2b). We observed two GFP-positive populations, most likely representing cells with mono- vs bi-allelic GFP insertions. We sorted the high-GFP population to generate polyclonal populations of cells with homogeneously high expression of GFP-CLTA. Similar results were observed when GFP was inserted at the N-terminus of the Rab11a gene (Extended Data Fig. 3a). These data demonstrate that Cas9EDVs can be combined with AAV-mediated delivery of HDR donor templates to fluorescently tag endogenous proteins in these myeloid cell lines, which may be a useful research strategy enabling cell biological investigations of innate immune signaling pathways.

**Figure 2:**
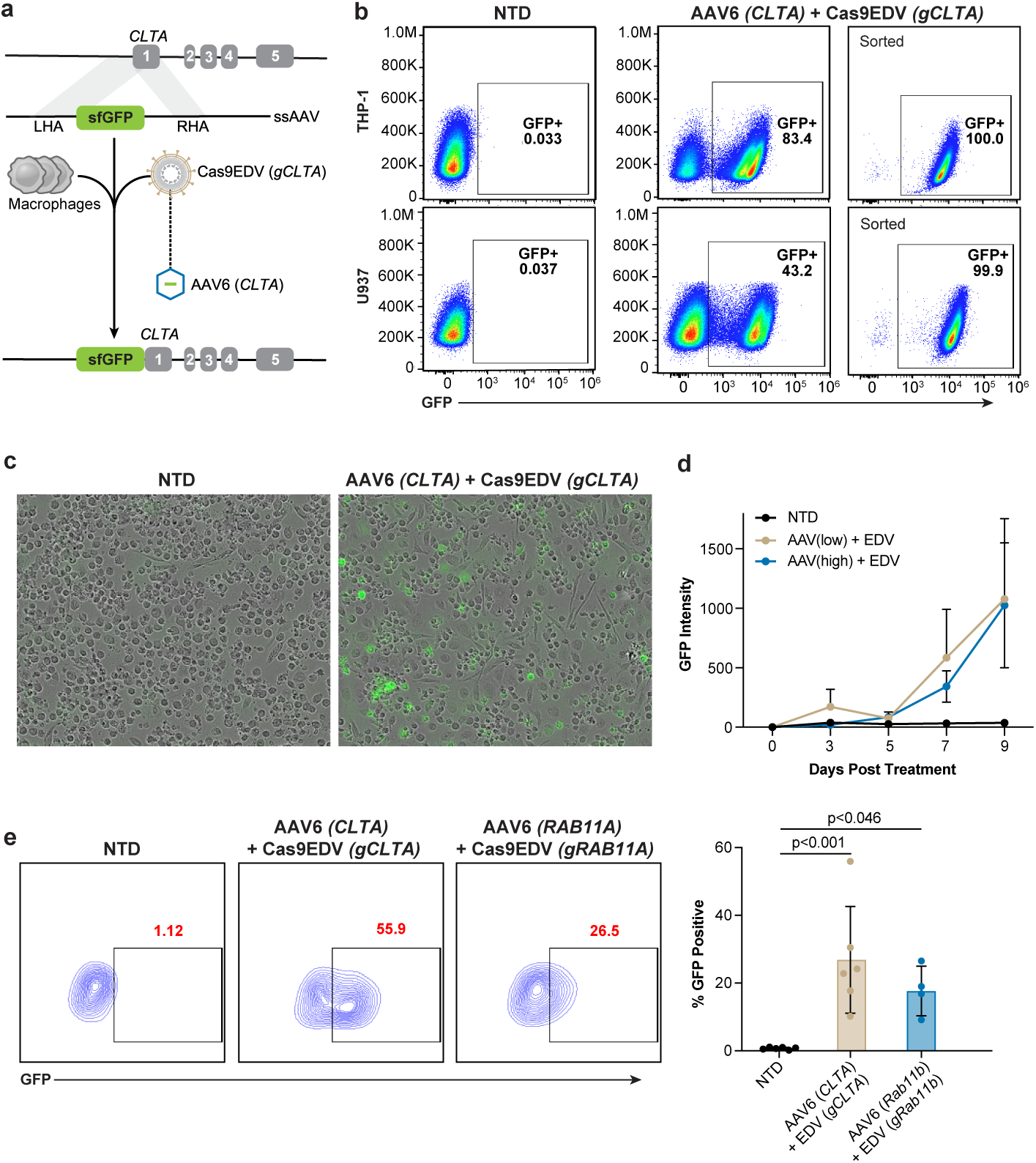
Cas9EDV/AAV co-treatment allows for site-specific knock-in of DNA sequences into the genome of human macrophages. **a**, Schematic representation of knock-in strategy, exemplified by the insertion of GFP into the N-terminus of CLTA. DNA donor template for homology mediated repair comprising the sequence encoding GFP together with left and right homology arms (LHA, RLA) is packaged into AAV6 particles. Macrophages are co-treated with these AAV particles and Cas9EDVs targeted near the start codon of the endogenous CLTA gene, so that successful knock-in should result in the production of a GFP-CLTA fusion protein. **b**, Flow cytometry analysis of THP-1 and U936 cells treated with CLTA-targeted Cas9EDV and AAV6 encoding donor template for insertion of GFP. Successful knock-in is read out by the appearance of GFP fluorescence. Cells were sorted to yield stable cell lines with homogenous GFP-CLTA expression. **c-e**, Live cell-imaging analysis (**c,d**) and Flow cytometry (**e**) of human primary macrophages (differentiated using GM-CSF) treated with CLTA-targeted Cas9EDV and AAV6 encoding donor template for insertion of GFP on Day 4 post-isolation. Successful knock-in is read out by the appearance of GFP fluorescence by live cell imaging on an incucyte instrument at indicated time points (**d**) or by flow cytometry on Day 9 post-Cas9EDV/AAV treatment (**e**). *P* values calculated by means of Dunnett’s multiple comparison test after ordinary one-way ANOVA tests.

We then applied this strategy to primary human macrophages. Macrophages were treated with HDR donor AAVs and Cas9EDVs targeting CLTA or Rab11a on Day 4 post isolation and GFP fluorescence was assessed by live-cell imaging and flow cytometry. Monitoring GFP expression by live-cell imaging over time revealed that fluorescent signals became first detectable at 7 days post-treatment (Fig. 2c, d). Flow cytometry analysis performed 13 days post-treatment revealed GFP fluorescence in up to approximately 55% and 25% of cells treated with CLTA- or Rab11a-specific AAV6/Cas9EDV combinations, respectively, indicating successful insertion of the GFP coding sequence into the genome of a substantial fraction of cells (Fig. 2e). Together, these results demonstrate that promoter-less transgene knock-in using AAV6 donors in combination with Cas9EDV enables precise, site-specific integration in human myeloid cells.

### VLP-mediated delivery enables base editing and epigenetic silencing of target genes in primary human macrophages

The design of additional eVLP systems have been reported for the delivery of gene editing enzymes beyond wildtype Cas9, including CRISPR-based base editors (BEs), which can introduce specific point mutations in the genome by converting adenine bases to guanine (adenine base editor; ABE) and/or cytosine to thymine (cytosine base editor; CBE)^29,30,42,43^. We treated human macrophages with eVLPs containing the ABE8e or the TadCBEd enzymes loaded with an sgRNA targeting a conserved splice site in *B2M*. Successful introduction of point mutations at this site is expected to disrupt production of B2M protein by interfering with proper production of mature mRNAs. Indeed, treatment of macrophages with ABE8e and TadCBEd-eVLPs resulted in a dose-dependent reduction of B2M protein on the cell surface, consistent with the corresponding rates of base conversions observed at the genomic level (Fig. 3a-c). Both ABE8e-eVLPs and TadCBEd-eVLPs achieved up to around 85% base editing in the expected window. Unexpectedly, in macrophages treated with ABE8e-eVLPs, we observed a relatively high degree of C-to-T conversion in addition to the expected A-to-G edits within the predicted editing window (Extended Data Fig. 4a-c). This bystander editing was only noted in the case of the ABE8e-eVLPs, and was not observed in the TadCBEd-eVLPs. It was previously reported that ABE enzymes have the capacity to also catalyze C-to-T conversions, albeit to low levels and only in specific sequence contexts^44^. Hence, these findings may hint at differences in editing biochemistries or DNA repair pathways in myeloid cells relative to other cell types. Nonetheless, eVLPs now offer an effective approach to introduce base edits into primary human myeloid cells.

**Figure 3:**
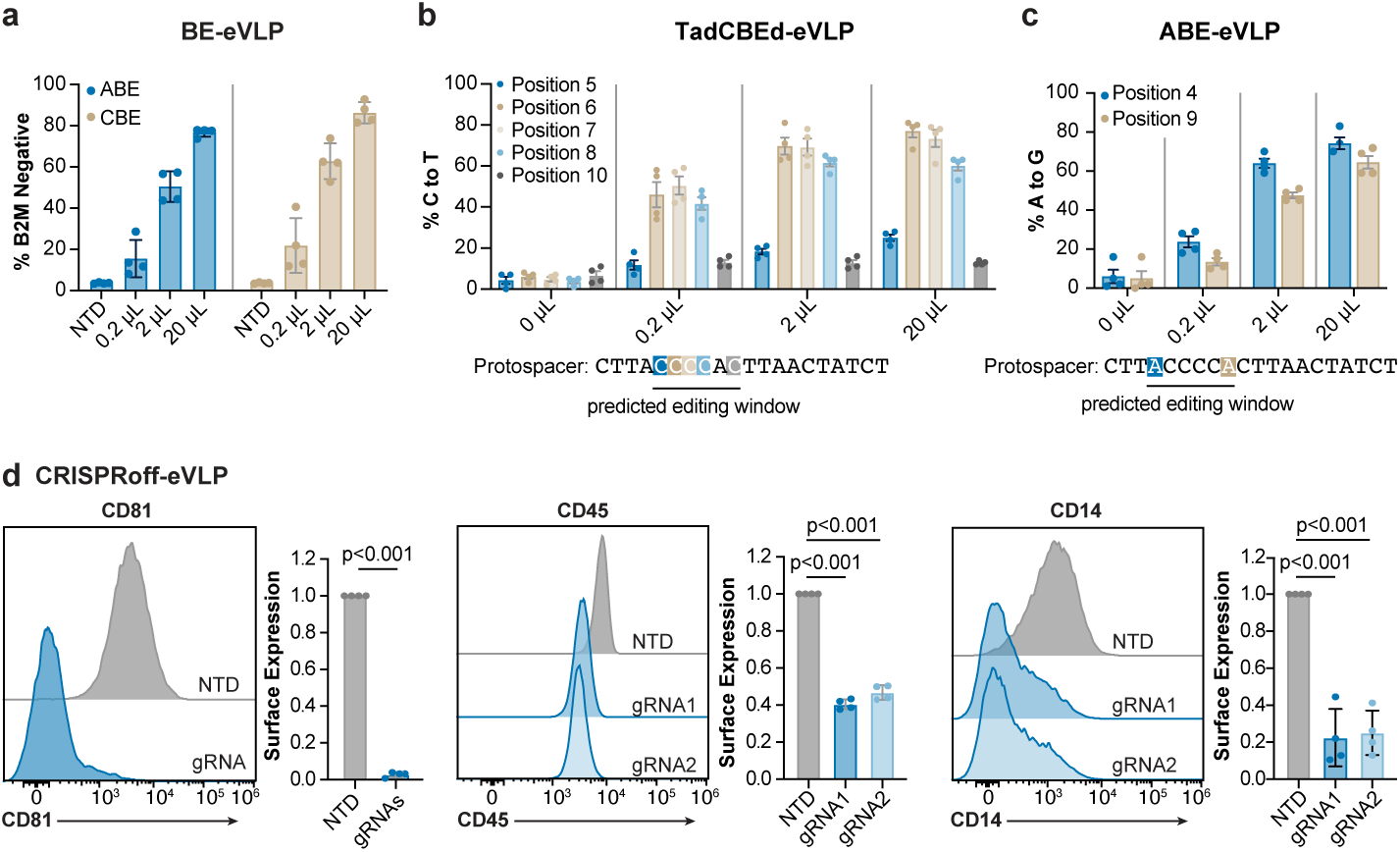
eVLPs enable base editing and epigenetic silencing of target genes in human macrophages. **a**, Quantification of B2M-deficient fraction of cells after treatment of macrophages with ABE8e-eVLPs or TadCBEd-eVLPs loaded with a sgRNA targeting a splice site in the *B2M* gene as determined by flow cytometry. **b**, Quantification of base editing after treatment of macrophages with TadCBEd-eVLPs loaded with a sgRNA targeting a splice site in the *B2M* gene. **c**, Quantification of base editing after treatment of macrophages with ABE8e-eVLPs loaded with a sgRNA targeting a splice site in the *B2M* gene. **d**, Quantification of CD81, CD45 or CD14 surface expression as determined by flow cytometry after treatment of macrophages with CRISPRoff-eVLPs targeting transcriptional start sites of the respective gene. Surface expression is displayed as median fluorescence intensity normalized to non-treated controls (NTD). *P* values calculated by means of Dunnett’s multiple comparison test after ordinary one-way ANOVA tests. Each data point represents the result of one independent assay. Bars and error bars represent mean ± standard deviation. Flow plots are representative of at least three independent experiments.

CRISPR nuclease and base editing both introduce permanent genetic alterations into target cells. Cas9 can also be engineered to reprogram epigenetic states at target loci without any changes to the DNA sequence, which enables a broad set of research and potential therapeutic applications^45^. eVLPs were recently engineered to deliver a catalytically inactive Cas9 (dCas9) fused to the Zim3 KRAB domain, DNA methyltransferase 3A (DNMT3A), and DNA methyltransferase 3-like protein (DNMT3L), enabling stable epigenetic silencing of target genes^46^. We used CRISPRoff-eVLPs loaded with sgRNAs targeting the transcriptional start sites of selected genes to silence expression of the tetraspanin CD81, as well as the myeloid-specific cell surface markers CD14 and CD45 (Fig. 3d). Taken together, these data demonstrate that eVLP-based RNP delivery systems enable high efficiency genome and epigenome editing in human primary myeloid cells.

### SLICeVLP enables large-scale pooled screening and identification of regulators of TNF production in primary macrophages

Having demonstrated the utility of VLP-mediated RNP delivery for targeting of individual genes in human primary myeloid cells, we investigated whether this technology may be adapted for high-throughput pooled CRISPR screening. Previously established platforms for genetic screening in human hematopoietic cells rely on the delivery of Cas9 protein via electroporation or introduction of Cas9-expression cassettes via lentiviral transduction^47–49^, which in our hands led to high toxicity, cell dysfunction, and/or low transduction and editing efficiency in primary myeloid cells. We attempted to deliver both a *B2M*-targeting sgRNA and Cas9-expression cassette into human macrophages by sequential transduction with a modified lentiviral vector that packages the Vpx protein of SIV-2 (LentiVPX), an antagonist of the myeloid-specific antiviral restriction factor SAMHD1 (Extended Data Fig. 5a, b)^50,51^. However, even though we were able to achieve relatively robust co-transduction, editing efficiency remained low (Extended Data Fig. 5c). Furthermore, the lentiVPX vector encoding Cas9 exhibited pronounced toxicity in macrophages at higher doses, thereby preventing large-scale pooled screening.

To address these challenges, we devised a system in which an sgRNA cassette is introduced via a LentiVPX vector, subsequently followed by Cas9 delivery by EDV or eVLP (Extended Data Fig. 5d). As a proof of concept, we first tested this strategy by targeting the B2M gene in human primary macrophages. On day 4 of differentiation, macrophages were transduced with a LentiVPX vector encoding a *B2M*-specific sgRNA as well as a BFP fluorescent marker, followed by treatment with sgRNA-empty EDV particles 24 hours later. This approach depleted cell-surface B2M protein in a modest fraction of BFP+ cells (Extended Data Fig. 5e). To improve editing efficiency, we generated EDVs packaging a non-targeting sgRNA (gScramble), reported to stabilize the Cas9 protein and enable sgRNA-swapping during Cas9 delivery^52^. This modification increased knockout efficiency by approximately two-fold compared to Cas9-only particles, consistent with previous reports using similar sgRNA-swapping strategies during Cas9 electroporation (Extended Data Fig. 5e, f)^52^. Furthermore, in this context, MMLV-based eVLP particles exhibited far superior editing efficiencies compared to the HIV-based EDV system, resulting in B2M depletion in >80% of BFP+ cells (Extended Data Fig. 5g). This high editing efficiency may be attributed to the enhanced capacity of v5 eVLPs for packaging large RNP complexes^30^. Taken together, we show that <u>s</u>gRNA lentiviral infection with <u>C</u>as9 delivery by <u>eVLP</u> (SLICeVLP) enables efficient gene disruption with traceable integrated sgRNA barcodes in primary human macrophages.

We next explored whether the SLICeVLP platform could be applied for large-scale loss-of-function pooled screening in human primary myeloid cells. To explore this, we first generated a library of sgRNA plasmids targeting 618 genes that are highly expressed in myeloid cells and have been implicated in innate immune signaling or identified as risk factors for various inflammatory diseases^53^. This library contained four sgRNAs per gene along with an additional 593 non-targeting sgRNAs, for a total of 3065 sgRNAs (Fig. 4a). For high-throughput screening, freshly isolated monocytes from the blood of two healthy human donors were differentiated into macrophages, transduced with a pool of LentiVPX vectors encoding this library, and then treated with gScramble-Cas9-eVLPs. 5 days after Cas9 delivery, cells were stimulated with LPS for 5 hours, fixed, and stained with a fluorescently labeled antibody against TNF. Cells were then sorted into bins of high and low TNF expressing cells (Top and bottom 20% of TNF intracellular expression level) and enrichment of sgRNAs in each sort bin was quantified by next-generation sequencing (Extended Data Fig. 6a). This screen revealed numerous positive and negative regulators of TNF production with high correlation between the two biological replicates performed in cells from different blood donors (Fig. 4b, Extended Data Fig. 6b and Supplementary Table 1). As expected, the *TNF* gene was identified as the top-ranking gene required for TNF protein production and there was no significant overall enrichment of non-targeting controls. Our screen identified additional known positive regulators of TNF production, including genes involved in extracellular LPS sensing (*TLR4, CD14, TICAM1, TICAM2, IRAK1)*^54^, NF-κB signaling (*NFKBIZ, NFKBIA, RELA*), MAP kinase signaling (*MAP3K8, MAP2K3*), as well as a number of transcription factors involved in inflammatory gene expression and macrophage differentiation (*IRF8, IRF1, MAF, IKZF1*^55,56^). On the other hand, we found that TNF production could be enhanced by targeting genes such as *TNFAIP3* (encoding the A20 protein, a known negative regulator of NF-κB signaling^57^), *ZFP36* and *R3CH1* (known to regulate mRNA stability)^4^, the transcription factors *MITF* and *FOS*, and *DUSP1*, a protein phosphatase with a reported role as a negative regulator of MAP kinase signaling^58^ (Extended Data Fig. 6c). To validate our pooled screen results, we treated macrophages with eVLPs carrying individual Cas9 RNPs coupled with sgRNAs targeting select hit genes, stimulated targeted cells with LPS, and assessed TNF production by flow cytometry. The results of these arrayed validation experiments were largely in concordance with the results of our pooled screen, although some hits with smaller, albeit consistent, effect sizes did not reach statistical significance (Fig. 4c, d and Extended Data Fig. 6d-f). Taken together, these data demonstrate that pooled CRISPR screens using SLICeVLP can be used to discover positive and negative regulators of human macrophage function.

**Figure 4:**
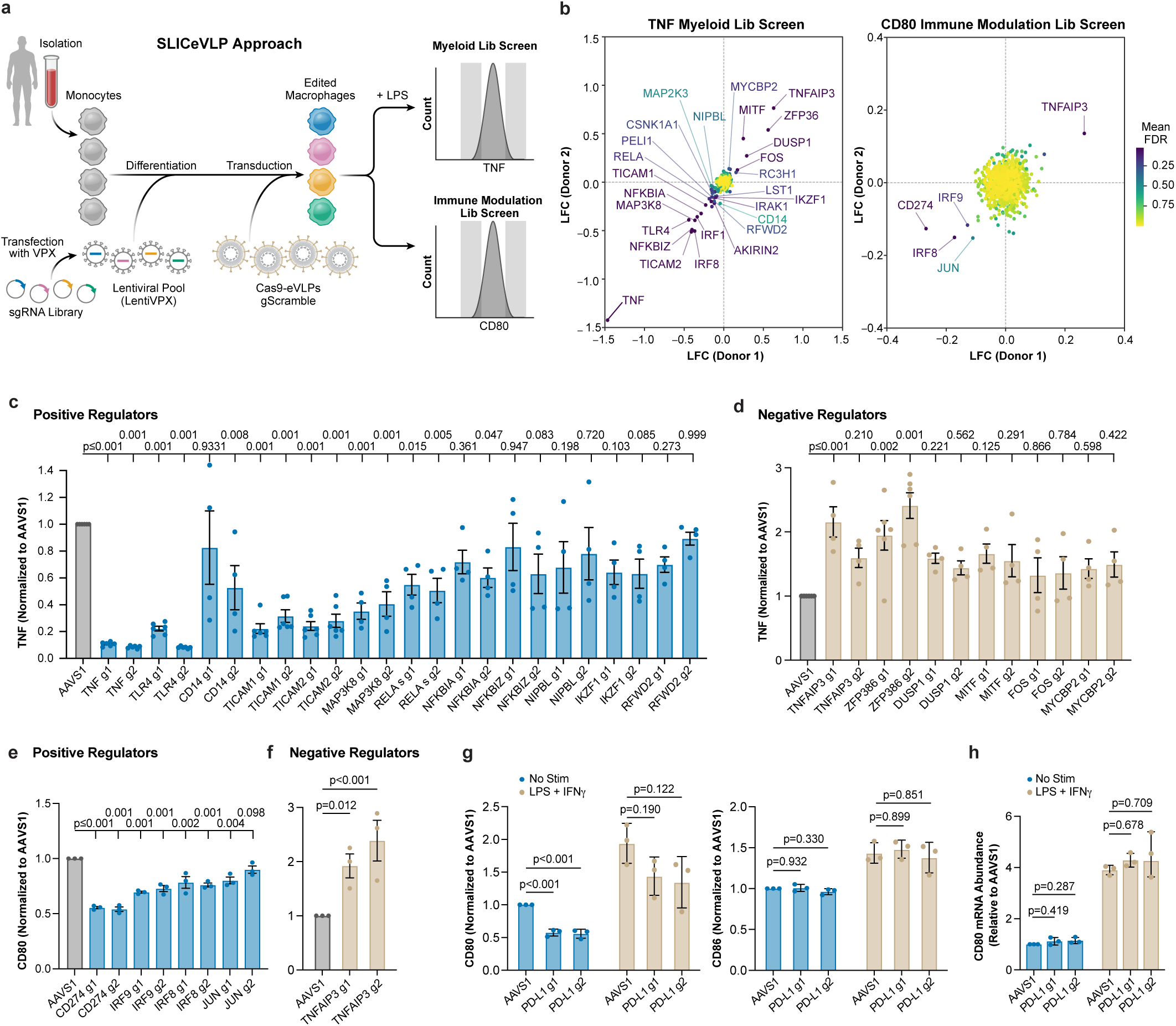
Pooled CRISPR loss-of-function screens in human primary macrophages identify regulators of TNF production and CD80 surface expression. **a**, Schematic of SLICeVLP strategy for pooled screening in human primary myeloid cells. **b**, Scatter plot showing correlation between two biological replicates of TNF production screen (left). X and Y axis show log-fold change for each individual screen replicate, and dots and gene names are colored based on the mean false discovery rate (FDR) between the two screens as calculated by MAGeCK. Scatter plot showing correlation between two biological replicates of screen for CD80 cell surface expression (right). X and Y axis show log-fold change for each individual screen replicate, and dots and gene names are colored based on the mean false discovery rate (FDR) between the two screens as calculated by MAGeCK. **c**,**d**, Arrayed validation of hits from TNF screen. Macrophages were treated with Cas9-eVLPs loaded with indicated sgRNAs, followed by stimulation with 50 ng/ml of LPS for 5 h and assessment of TNF production by intracellular cytokine staining and flow cytometry. Data is normalized to AAVS1-targeted control cells from each independent replicate. **e**,**f**, Arrayed validation of hits from TNF screen. Macrophages were treated with Cas9-eVLPs loaded with indicated sgRNAs, followed by assessment of CD80 surface levels by flow cytometry. Data is normalized to AAVS1-targeted control cells from each independent replicate. **g**, Macrophages were treated with Cas9-eVLPs loaded with sgRNAs targeting CD274/PD-L1 and stimulated with LPS/IFN-γ for 24 h or left non-treated, followed by assessment of CD80 and CD86 surface levels by flow cytometry. **h**, Macrophages were treated with Cas9-eVLPs loaded with sgRNAs targeting CD274/PD-L1 and stimulated with LPS/IFN-γ for 24 h or left non-treated, followed by assessment of CD80 mRNA abundance by quantitative RT-PCR. **g**,**h**, *P* values calculated by means of Tukey’s multiple comparison test after ordinary one-way ANOVA tests. Each data point represents the result of one independent assay. Bars and error bars represent mean ± standard deviation.

### Pooled screen identifies regulators of CD80 surface expression

Next, we employed SLICeVLP to perform a screen to identify regulators controlling surface expression of CD80, a potent co-stimulatory signal to T cells and marker of a pro-inflammatory macrophage state. The screen design was intended to focus specifically on factors that influence CD80 surface expression, and by extension, macrophage cell state, in the absence of any additional pro-inflammatory stimulus. We introduced a lentiviral sgRNA library targeting 1349 transcription factors and other known immunomodulatory molecules (4 sgRNAs per gene; 13 GFP-targeting and 593 non-targeting sgRNAs as controls; 6002 sgRNAs total)^59^ into macrophages freshly isolated from two healthy human donors, followed by Cas9-eVLP-mediated gene perturbation. After five additional days of culture, we stained and sorted for high and low CD80 surface expression (Top and bottom 20%) and quantified sgRNA abundance in each sorted bin (Extended Data Fig. 7a). Although the two donor replicates showed strong correlation, this screen identified only a small number of candidate hits, which were subsequently validated in an arrayed format (Fig. 4b, Extended Data Fig. 7b and Supplementary Table 2). Notably, akin to our TNF screen described above, *TNFAIP3* again emerged as the top negative regulator. Furthermore, we found that targeting the transcription factors, *IRF8, IRF9, and JUN* decreased CD80 surface levels, likely by interfering with macrophage differentiation or by dampening basal or eVLP-induced interferon signaling (Fig. 4e, f and Extended Data Fig. 7c, d).

Interestingly, we observed the strongest decrease in CD80 surface expression when targeting the gene *CD274*, encoding the protein PD-L1. This protein is most well-known for its role in regulating signaling by the immune checkpoint PD-1 on T cells. PD-L1 is only expressed at low levels in macrophages at baseline, but becomes upregulated upon stimulation with a cocktail of LPS and IFN-γ (Extended Data Fig. 8a). Using *CD274*-specific eVLPs, we were able to ablate PD-L1 surface expression in more than 95% of the targeted macrophage cells (Extended Data Fig. 8b). In line with our pooled screen results, perturbing PD-L1 in macrophages led to significant decreases in CD80 surface expression in both the presence and absence of additional stimulation with LPS and IFN-γ (Fig. 4g). Surface levels of the related costimulatory molecule CD86 as well as cytokine release in response to LPS/IFN-γ treatment were unaffected by PD-L1 knockout, indicating a CD80-specific effect rather than a changed overall cell state. Additionally, we observed that lower CD80 surface expression in PD-L1-deficient macrophages was accompanied by decreased total CD80 protein in whole-cell lysates, yet the abundance of CD80 mRNA was unchanged in PD-L1-deficient cells compared to *AAVS1*-targeted control (Fig. 4h and Extended Data Fig. 8c). These findings indicate a tightly coupled regulatory mechanism of CD80 by PD-L1 on the protein level. Indeed, previous studies have reported that CD80 and PD-L1 can interact in cis on the surface of cells and this interaction can inhibit binding of PD-L1 to its canonical binding partner PD-1 on neighboring cells in trans^60–62^. Thus, these collective data imply that binding of PD-L1 to CD80 may stabilize the latter on the cell surface and protect it from degradation, revealing complex cross talk between inhibitory and costimulatory signaling from myeloid cells to T cells.

### *TNFAIP3* ablation reprograms macrophages toward a pro-inflammatory state and increases HER2CAR macrophage cytotoxicity

Intrigued that both functional screens converged on *TNFAIP3* as the strongest negative regulator, we next investigated the functional consequences of *TNFAIP3* disruption in primary human macrophages. Macrophages were treated with Cas9-eVLPs carrying an sgRNA targeting *TNFAIP3*. Efficient knockout of *TNFAIP3* was validated at the genomic level and protein depletion was confirmed by immunoblot (Extended Data Fig. 9a, b). To further characterize the phenotype of *TNFAIP3*-deficient macrophages, we assessed CD80 and CD206 expression, well-established markers of M1 and M2 polarization, after stimulation with LPS/IFN-γ or IL-4, respectively. Ablation of *TNFAIP3* significantly increased expression of the M1 marker, CD80, and prevented the IL-4-induced upregulation of the M2 marker CD206 (Fig. 5a-d). Moreover, we observed production of inflammatory cytokines like TNF and IL-6 by *TNFAIP3*-deleted macrophages even in the absence of any additional stimulation (Fig. 5e). These data buttress the role of *TNFAIP3* as a master regulator controlling macrophage polarization and inflammatory status.

**Figure 5:**
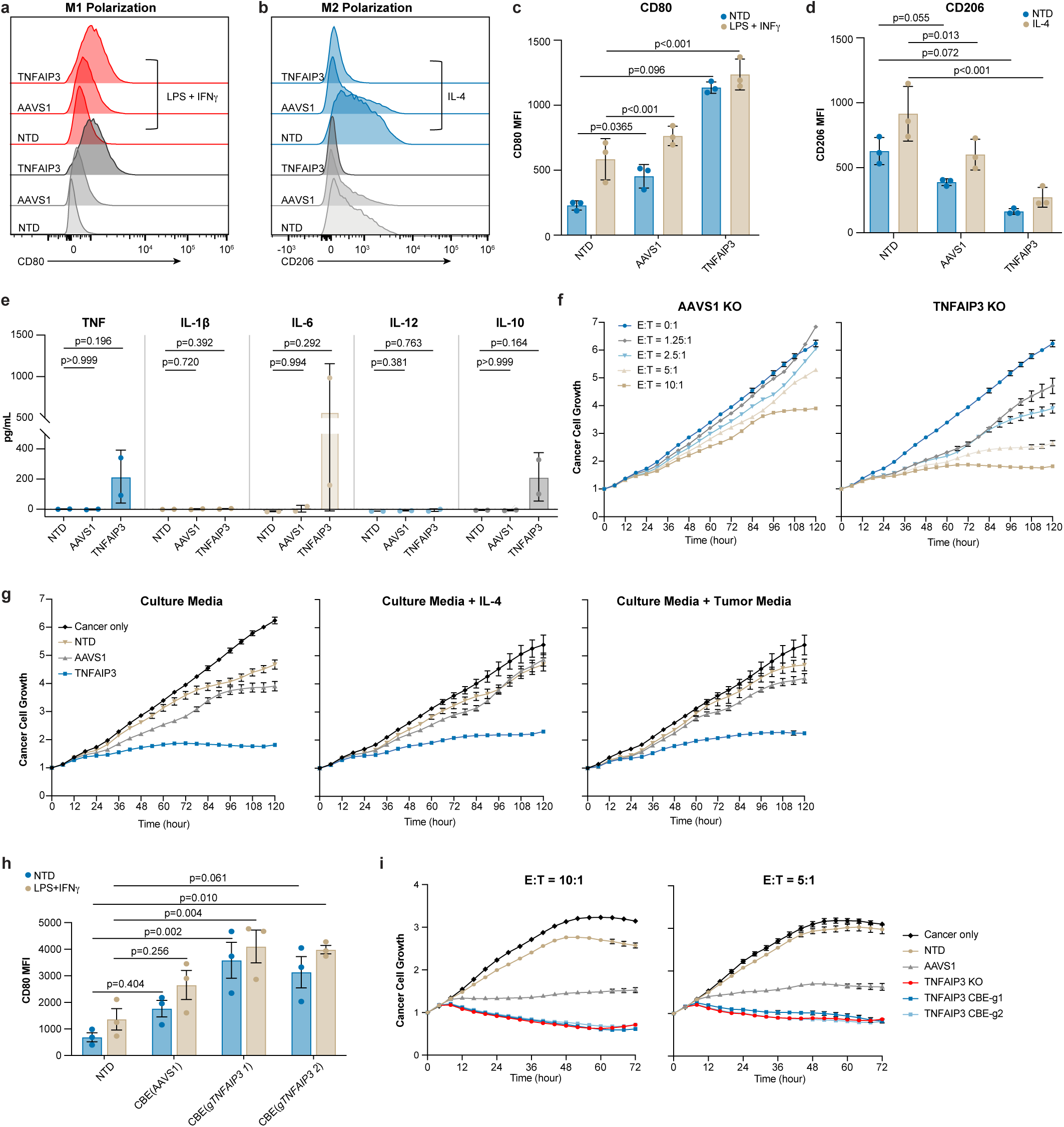
*TNFAIP3*-deficiency enhances tumor cell killing in a HER2-CAR macrophage model **a**,**b**, Macrophages were treated with Cas9-eVLPs loaded with sgRNAs targeting TNFAIP3 and stimulated with LPS/IFN-γ (**a**) or IL-4 (**b**) or left non-treated, followed by assessment of the M1 marker CD80 or the M2 marker CD206 by flow cytometry. **c**,**d**, Quantification of CD80 or CD206 expression in macrophages stimulated with LPS/IFN-γ (**c**) or IL-4 (**d**). *P* values calculated by means of Tukey’s multiple comparison test after ordinary two-way ANOVA tests. **e**, Analysis of cytokine release from macrophages treated with Cas9-eVLPs loaded with sgRNAs targeting TNFAIP3. *P* values calculated by means of Dunnett’s multiple comparison test after ordinary one-way ANOVA tests. **f**, Macrophages were engineered to express a HER2-specific CAR and treated with Cas9-eVLPs loaded with sgRNAs targeting *TNFAIP3* or *AAVS1*. Macrophages were then co-cultured with SK-OV3 cells stably expressing mKate at indicated Effector:tumor cell (E:T) ratios and tumor cell killing was assessed over time. **g,** Macrophages were engineered to express a HER2-specific CAR and treated with Cas9-eVLPs loaded with sgRNAs targeting TNFAIP3 or AAVS1. Macrophages were then co-cultured with SK-OV3 cells stably expressing mKate at an E:T ratio of 5:1 in the presence of indicated immunosuppressive stimuli and tumor cell killing was assessed over time. **h**, Macrophages were treated with CBE-eVLPs loaded with sgRNAs designed to introduce pre-mature stop-codons in the TNFAIP3 gene and stimulated with LPS/IFN-γ or left non-treated, followed by assessment of CD80 surface levels by flow cytometry. *P* values calculated by means of Tukey’s multiple comparison test after ordinary two-way ANOVA tests. **i**, Macrophages were engineered to express a HER2-specific CAR and treated with Cas9-eVLPs or CBE-eVLPs loaded with sgRNAs targeting TNFAIP3. Macrophages were then co-cultured with SK-OV3 cells stably expressing mKate at indicated E:T ratios and tumor cell killing was assessed over time. Each data point represents the result of one independent assay. Bars and error bars represent mean ± standard deviation. Flow cytometry plots and tumor cell growth curves are representative of at least three independent experiments.

Since *TNFAIP3* ablation drives macrophages toward an inflammatory state, we explored whether it may also enhance the anti-cancer potential of engineered cells in a CAR macrophage context. Leveraging the LentiVPX system, we generated macrophages expressing a HER2-specific chimeric antigen receptor (HER2CAR) bearing a CD3zeta signaling domain (Extended Data Fig. 9c). We then co-cultured HER2CAR-Macrophages with SK-OV3 ovarian cancer cells stably overexpressing mKate and assessed tumor cell growth by live-cell imaging. Strikingly, we observed that *TNFAIP3*-deleted HER2CAR macrophages exhibited significantly enhanced anti-tumor activity against SK-OV3 tumor cells compared to AAVS1-targeted control cells expressing the same CAR. This phenotype was consistent across multiple donors and multiple tested effector-to-target ratios (Fig.5f) and was also evident under immunosuppressive conditions (Fig. 5g). We tested whether similar enhancement of CAR macrophages could be achieved with base editing instead of Cas9 nuclease, which would avoid liabilities related to double-stranded DNA breaks. To this end, we designed two sgRNAs to introduce a premature stop codon in the *TNFAIP3* gene when co-delivered with a CBE enzyme. Indeed, treatment of primary human macrophages with TadCBEd-eVLPs carrying these sgRNAs led to the efficient depletion of A20(*TNFAIP3*) protein (Extended Data Fig. 9d) and polarized cells towards an inflammatory M1 fate, as indicated by upregulation of CD80 (Fig. 5h). TadCBEd-eVLP-mediated introduction of stop codons in *TNFAIP3* also enhanced the tumoricidal capacities of HER2CAR-macrophages, to a similar extent as disruption of *TNFAIP3* using Cas9 nuclease-eVLPs (Fig. 5i). Taken together, these findings indicate that gene editing of key regulators such as *TNFAIP3* - either by Cas9 nuclease-mediated gene disruption or by base editing - can enhance M1 polarization, cause resistance to M2 polarizing cues, and improve CAR-macrophage anti-cancer function.

## Discussion

In this study, we present a technology platform for high-efficiency genome engineering in primary human myeloid cells, overcoming long-standing barriers that have limited both mechanistic studies and translational applications in this clinically important cell lineage. By leveraging VLP-mediated delivery of diverse CRISPR editing enzymes, we achieved robust gene knockout, base editing, epigenetic silencing, and site-specific knock-in in primary monocytes, macrophages, and dendritic cells. A major advantage of VLP-mediated genome editing over previously described nucleofection-based methods is the preservation of cell viability and functional responsiveness to innate immune stimuli like LPS and IFN-γ. Bulk RNA-seq studies revealed that nucleofection triggers broad transcriptional reprogramming toward hyporesponsive and immunosuppressive macrophage phenotypes, whereas VLP-engineered cells retained transcriptional hallmarks of terminally differentiated macrophages and mounted intact inflammatory responses. While VLP treatment slightly skewed macrophages towards an inflammatory (‘M1’) phenotype, this feature may even be beneficial in contexts such as CAR macrophage therapy, where activation of interferon and pro-inflammatory signaling pathways correlates with improved anti-tumor activity^15,63^. This modest skew towards an inflammatory (‘M1’) phenotype is strongly preferable to the immunosuppressive and immune-unresponsive cell state that we observed following nucleofection.

The translational potential of this platform can be expanded through the development of cell type-specific envelopes, as demonstrated by the design of a CD11b-targeted fusogen that enables selective delivery of genetic payloads to myeloid cells. Given the emerging success of in vivo manufacturing approaches for T and NK cell therapies^28,40,41,64^, analogous strategies for myeloid cells could circumvent the limitations of ex vivo culture, including manufacturing complexity, high cost, and manipulation-induced dysfunction. The targeting approach using a target-specific scFv in conjunction with VSVGmut is inherently modular^36^, suggesting that envelope specificity could be further tuned to distinct myeloid subsets or activation states, offering a level of precision not currently achievable in myeloid-directed therapies.

The high efficiency and low toxicity of VLP-mediated Cas9 delivery enabled pooled genetic screens in monocyte-derived macrophages, representing, to our knowledge, the first large-scale pooled CRISPR-based functional genomics analyses in any primary human myeloid cell type. Developing technologies for pooled screening in human myeloid cells has been particularly challenging, in part because peripheral blood monocytes are generally considered post-mitotic and do not proliferate ex vivo. This severely limits the number of cells that can be obtained from each blood donation, and necessitates especially gentle genome editing approaches to preserve sufficient cell yield. The SLICeVLP workflow addresses these constraints, providing a flexible and versatile platform that can be adapted to interrogate diverse genetic programs governing primary myeloid cell biology. While our screens have focused on macrophages, this approach could be applied to other myeloid subsets, such as dendritic cells, for example, to investigate genetic determinants of antigen uptake by phagocytosis, antigen presentation, or responses to bacterial and viral pathogens. More broadly, SLICeVLP could open pooled genetic screening in other non-dividing cell types that are refractory to Cas9 delivery by electroporation or lentiviral transduction. Future studies should expand the functionality of SLICeVLP by incorporating more advanced editing approaches and readouts. For instance, BE-eVLPs may be used to systematically interrogate the functional effects of single nucleotide mutations^49^, and the coupling of targeted genetic perturbations to single-cell transcriptomics^53,65^ will allow for the mapping of gene regulatory networks in a more granular fashion than the FACS-based assays used herein. In addition, further optimization of eVLP architecture, delivery properties, and production strategies combined with improved cell sourcing and differentiation protocols may enable screening of substantially larger libraries and ultimately make genome-wide analyses in primary human myeloid cells feasible.

Our screens for regulators of TNF production and CD80 surface expression converged on *TNFAIP3* as a central negative regulator of pro-inflammatory macrophage states. Ablation of *TNFAIP3* induced M1 polarization, facilitated inflammatory cytokine production in the absence of additional stimuli, and rendered macrophages resistant to IL-4-driven M2 skewing. Collectively, these changes enhanced the tumoricidal activity of HER2CAR macrophages. These findings extend prior observations that *TNFAIP3* inhibits NF-κB signaling through control of ubiquitination and deubiquitination of key pathway components such as NEMO and RIPK1^66–68^. Notably, previous genetic screens in primary T cells identified *TNFAIP3* as a target to enhance activation of and cytokine production by T cells, and by extension CAR T cells, as well as the ability of these cells to mediate tumor cell killing^48,69^. However, the potential of TNFAIP3-deletion to enhance macrophage therapies had not been previously explored. This case study highlights the power of our genetic screening platform to identify and prioritize candidate gene modifications that increase the potency of myeloid cell-based immunotherapies.

In summary, our VLP-based engineering platform and SLICeVLP screening pipeline establish a foundation for both mechanistic discovery and therapeutic innovation in human myeloid biology. By combining efficient, non-toxic delivery of diverse CRISPR modalities with scalable pooled screening, this approach will enable systematic mapping of regulatory networks in myeloid cells and guide the rational design of next-generation myeloid cell therapies.

## Material and Methods

### Isolation of human primary monocytes, and differentiation into macrophages and dendritic cells

Primary human monocytes were isolated from fresh peripheral blood Leukopaks (STEMCELL Technologies) with institutional review board-approved informed written consent. Leukopaks were washed twice with EasySep buffer (sterile PBS pH 7.4, 2% FCS, 1 mM EDTA pH 8.0) before isolation of monocytes using the Human Monocyte Enrichment kit without CD16 depletion (STEMCELL Technologies) according to manufacturer’s instruction. Briefly, PBMCs were diluted to 100x10^6^ cells/ml in EasySep buffer, mixed with 100 µl of antibody cocktail per 100 million cells and incubated at RT for 5-10 mins. Magnetic beads were added and following 3 min of incubation at RT, tubes containing cells were placed in a magnetic field for 10 mins. Unbound cells (pure monocyte fraction) were removed, washed one more time with EasySep buffer. Cells were then counted and resuspended in Immunocult SF Macrophage media (STEMCELL technology) supplemented with appropriate cytokines at a density of 1x10^6^ cells/ml and seeded in TC-treated 24-well plate (for most experiments; 0.5x10^6^ cells per well) or 12-well plates (for pooled screens; 2x10^6^ cells per well). For macrophage differentiation, media was supplemented with 20 ng/ml of M-CSF or GM-CSF, for dendritic cell differentiation media was supplemented with 20 ng/ml of GM-CSF and 20 ng/ml of IL-4 (all from Peprotech). Media was replenished every 24 - 72 h by adding at least one half volume of fresh media and cytokines. All tissue culture was performed under sterile conditions and cells were cultured in temperature-controlled incubators at 37°C and 5% CO_2_.

### Production of lentiviral vectors

Vpx-containing lentiviral vectors were produced as described in Norton et. al.^51^ with slight modifications. LentiX-HEK293T packaging cells were cultured in complete Opti-MEM supplemented with 5% fetal calf serum at 37 °C in 5% CO₂. 11 × 10⁶ HEK293T cells were plated in a T-75 flask containing 15 mL of complete Opti-MEM and grown until approximately 90% confluence (18–24 h). On the day of transfection, transfer plasmid encoding the HER2-CAR or sgRNAs (10 µg), the packaging plasmid pMDL-chp6 (10 µg), the VSV-G envelope plasmid pMD2.G (3.5 µg), the Rev expression plasmid (2.5 µg), and the Vpx expression plasmid (1 µg) were combined in 2 ml of Opti-MEM. To this mixture, 56 µL of (P3000 reagent ThermoFisher) was added and gently mixed. Separately, 60.3 µL of Lipofectamine 3000 (ThermoFisher) was diluted in 1.9 mL of Opti-MEM. Lipofectamine solution was then combined with the DNA–P3000 mix and incubated at room temperature in a biosafety cabinet for 15 minutes to allow complex formation. Immediately prior to transfection, 7 mL of medium was removed from each flask, and approximately 4 mL of the DNA–Lipofectamine complex was added dropwise onto the cells. After incubation at 37 °C for 6 hours, the transfection mixture was aspirated and replaced with 15 mL of fresh complete Opti-MEM supplemented with ViralBoost (1:500; Alstem). Cells were then returned to the incubator and maintained for a total 48 hours post-transfection. At 24 and 48 hours post initial transfection, culture supernatants were collected and clarified by centrifugation at 300 × g for 5 minutes to remove cellular debris. Five milliliters of Lenti-X Concentrator (TakaraBio) was added to each supernatant, mixed by gentle inversion, and stored overnight at 4 °C. Virus–concentrator mixtures were centrifuged at 1,500 × g for 45 minutes at 4 °C; supernatants were caully discarded, and tubes were spun again at 1,500 × g for 1 minute to remove residual fluid. Viral pellets from both the first and, if performed, second harvest were resuspended together in 150 µL of Opti-MEM, aliquoted and stored at –80 °C. When larger amounts of viral vectors were required, transfections were performed in T225 flasks by tripling all volumes, plasmid and cell amounts listed above.

### Production of EDVs and eVLPs

Cas9EDVs, v5 Cas9-eVLPs, v5 ABE8e-eVLPs, v3b TadCBEd-eVLPs and CRISPRoff-eVLPs (targeting CD14 and CD45) were produced as described in Hamilton et al, 2024, Raguram et al 2024, An et al 2024, and Xu et al 2025, respectively, with modifications^28,30,43,46^. LentiX-HEK293T packaging cells were cultured in complete Opti-MEM supplemented with 5% fetal calf serum at 37 °C in 5% CO₂. 32 × 10⁶ HEK293T cells were plated in a T-225 flask containing 45 mL of complete Opti-MEM and grown until approximately 90% confluence (18–24 h). For Cas9EDV production, plasmids encoding HIV Gag-pol (15 µg), HIV Gag-Cas9 (30 µg) and VSV-G (3.2 µg) were mixed with 5.5 ml of Opti-MEM. Both Gag-Pol and Gag-Cas9 encoding plasmids also encoded the sgRNA targeting the gene of interest under control of the U6 promoter. For Cas9-eVLP production, plasmids encoding v5 MMLV Gag-pol (27 µg), v5 MMLV Gag-Cas9 (9 µg), VSV-G (3.2 µg) and an sgRNA under the control of a U6 promoter (35.2 µg) were combined in 5.5 ml of Opti-MEM. For ABE8e-eVLP production, v5 MMLV Gag-pol (27 µg), v5 MMLV Gag-ABE8e (9 µg), the VSV-G (3.2 µg) and an sgRNA under the control of a U6 promoter (35.2 µg) were combined in 5.5 ml of Opti-MEM. For TadCBE-eVLP production, plasmids encoding MMLV Gag-pol (22.5 µg), MMLV Gag-COM-Pol (16 µg), MMLV Gag-P3-Pol (3.38 µg), P4-TadCBEd (3.38 µg), VSV-G (3.2 µg) and an sgRNA containing the Com aptamer under the control of a U6 promoter (35.2 µg) were combined in 5.5 ml of Opti-MEM. For CRISPRoff-eVLP production, plasmids encoding MMLV Gag-pol (3.6 µg), MMLV Gag-Zim3-CRISPRoff (32.4 µg), VSV-G (4 µg) and an sgRNA under the control of a U6 promoter (40 µg) were combined in 5.5 ml of Opti-MEM. To these mixtures, 167 µL of P3000 reagent (ThermoFisher) was added and gently mixed. Separately, 181 µL of Lipofectamine 3000 (ThermoFisher) was diluted in 5.5 mL of Opti-MEM. Lipofectamine solution was then combined with the DNA–P3000 mix and incubated at room temperature for 15 min. Immediately prior to transfection, 20 mL of medium was removed from each flask, and approximately 11 mL of the DNA–Lipofectamine complex was added dropwise onto the cells. After incubation at 37 °C for 6 hours, the transfection mixture was aspirated and replaced with 40 mL of fresh complete Opti-MEM supplemented with ViralBoost (1:500; Alstem). Cells were then returned to the incubator and maintained for a total 48 hours. Cell culture supernatants were collected and clarified by centrifugation at 500 × g for 5 minutes to remove cellular debris. 15 ml of Lenti-X Concentrator (TakaraBio) was added to each supernatant, mixed by gentle inversion, and stored overnight at 4 °C. Virus–concentrator mixtures were centrifuged at 1,500 × g for 45 minutes at 4 °C; supernatants were carefully discarded, and tubes were spun again at 1,500 × g for 1 minute to remove residual fluid. Viral pellets were resuspended in Opti-MEM (1:100 of starting culture volume), aliquoted and stored at –80 °C. CRISPRoff-eVLPs carrying a combination of two sgRNAs targeting CD81 were produced exactly as described in Xu et al, 2025^46^.

### Lentiviral transduction and EDV/eVLP treatments

Unless indicated otherwise, EDV/eVLP treatments or lentiviral transductions of human primary myeloid cells were performed with cells plated in TC-treated 24-well plates. Indicated amounts of VLPs/lentiviral vectors were mixed with 500 µl of pre-warmed SF macrophage media supplemented with appropriate cytokines. Media was aspirated from cells and EDV/eVLP-containing media was added followed by spinfection at 1250 g and 32°C for 1 h. Cells were incubated overnight before swapping media and returning to routine culture. Assessment of successful gene disruption was performed after 4 - 5 days for experiments with Cas9-eVLPs and BE-eVLPs, and 8 days for CRISPRoff-eVLPs. For arrayed validation experiments for pooled screens, macrophages were treated with 20 µl of Cas9-eVLPs targeting indicated the gene per well on Day 3 or 4 post-isolation and analyzed or further stimulated after 5 days of incubation. For the arrayed validation of the TNFAIP3 hit in the TNF screen, only data points from cells treated with eVLPs on Day 4 (not Day 3) were included.

For cell yield analyses, cells from one well per treatment conditions were lifted using accutase, stained with Trypan Blue (1:1 v/v ratio) and numbers of viable cell was (Trypan Blue negative) assessed on a Countess Automated Cell Counting Device (Thermo Fisher Scientific). Relative cell yield was calculated by normalizing to the viable cell counts obtained from non-treated control wells containing cells from the same donor.

### Analysis of VLP-treated cells by flow cytometry

Following treatment of cells with EDVs or eVLPs, media was aspirated from cells and 0.5 ml of accutase solution (BD) was added to each well of a 24-well plate. After incubation at 37 °C for 30 min, cells were rinsed off by pipetting and collected into a U-bottom 96-well plate for flow cytometry staining. For surface staining, cells were washed once (400 g, 5 min) with 200 μl of EasySep buffer before staining with human Trustain FC-receptor block (1:200; Biolegend), Ghostdye Red780 or Violet450 (1:1000; Tonbo) and one of the following antibodies in EasySep buffer for 30 min on ice in the dark: PacificBlue anti-B2M, Alexa657 anti-CD45, APC anti-CD81, APC anti-CD14, APC anti-CD86, APC or PE anti-CD80, APC anti-CD274/PD-L1 (all from Biolegend and used at 1:200 dilution). Cells were washed once in 200 μl EasySep buffer before analysis on the Thermo Fisher Attune NxT flow cytometer with autosampler (A29004).

For staining of intracellular TNF, cells were transduced with indicated eVLPs as described above and after 5 days, cells were stimulated in a 24-well plate 500 μl of SF macrophage media containing 50 ng/ml of ultrapure LPS (Enzo Life sciences) and GolgiPlug reagent (1:1000; BD) for 5 h. Cells were then lifted with 500 µl of accutase and 200 – 300 µl of cell suspension was collected into a U-bottom 96-well plate. Cells were washed once (400 g, 5 min) with 200 μl of EasySep buffer before staining with human Trustain FC-receptor block (1:200; Biolegend) Ghostdye Violet450 (1:1000; Tonbo) for 20 min on ice in the dark. Cells were then fixed with 200 μl of Cytofix reagent (BD Biosciences) for 30 min on ice, washed once with 200 μl of Perm/Wash buffer and stained with APC-conjugated anti-TNF antibody (Biolegend) diluted 1:200 in Perm/Wash buffer for 30 min on ice in the dark. Cells were washed once in Perm/Wash buffer before analysis on the Thermo Fisher Attune NxT flow cytometer.

### Nucleofection of Cas9 ribonucleoproteins (RNPs) into primary human monocytes

For RNP assembly, lyophilized crRNA and tracrRNA (Dharmacon) were reconstituted to 160 µM in Nuclease-Free Duplex buffer (IDT), and used immediately or aliquoted and stored at –80 °C. Equal volumes of crRNA and tracrRNA were mixed and incubated at 37 °C for 30 min to form into an 80 µM gRNA duplex. The duplex was then combined 1:1 (vol/vol) with 40 µM SpCas9 protein (UC Macrolab) to yield a 2:1 RNA:Cas9 molar ratio and incubated at 37 °C for 15 min, producing 20 µM Cas9 RNP. RNPs were used immediately or stored at –80 °C. Purified CD14⁺ monocytes (1∼2e6 cells per reaction) were resuspended in 20 µL room-temperature P3 nucleofection buffer (Lonza). 2.5 µL of the appropriate Cas9 RNP was added per reaction and the cell/RNP mixture was gently mixed and transferred to a 96-well nucleofection cuvette plate compatible with the Lonza 4D X unit or Shuttle unit (Lonza). Nucleofection was performed using program CM-137. Immediately after pulsing, 80 µL of pre-warmed culture medium was added directly to each cuvette, and cells were transferred to culture plates pre-filled with the appropriate medium for differentiation and subsequent analyses.

### Bulk RNA seq analysis

Isolated CD14⁺ monocytes were subjected to nucleofection or Cas9EDV-mediated gene editing targeting the *AAVS1* locus on day 0. Monocyte samples were collected 1 day post-editing, and macrophage samples were harvested 7 days post-editing. Cells were washed twice with DPBS, pelleted by centrifugation. RNA extraction, library preparation, and next-generation sequencing was performed by GENEWIZ Inc. For each pair of differential expression analyses, we ranked the genes based on their Wald statistics. We then used the fgsea function in the fgsea package (version 1.34.2) to calculate their enrichment in hallmark gene sets from the MsigDb database^70^. To evaluate the expression of macrophage or monocyte subtype-specific marker genes in each condition, we calculated the enrichment score of each gene set in each sample using GSVA software (version 2.2.0). We utilized the variance-stabilizing transformed expression data for each sample and ran GSVA with the kcdf = “Gaussian” option. As gene sets, we utilized the marker genes for monocyte or macrophage subtypes defined in single-cell RNA-seq analysis^34^. For the CD14CTX marker^31^, we used the reported comparison table of CD14CTX vs. CD14APC gene expression and selected the top 300 significant genes with a log fold change > 0.3

### Genotyping

To analyze editing outcomes on the genome level, ∼0.1x10^6^ cells were collected in a PCR strip tube and centrifuged in a table top minifuge. Cell pellets were resuspended in 50 μl Quickextract solution (Lucigen; QE09050) and genomic DNA was extracted by incubation at 65°C for 6 min, followed by heat inactivation at 95°C for 2 min. Genomic loci of interest were amplified by touchdown PCR using 2X KAPA HiFi master mix (Roche) or Phusion Hotstart polymerase (NEB). Amplicons were purified by column purification using the Nucleospin Gel and PCR clean up kit (Takara Bio) according to manufacturer’s protocols and analyzed by sanger sequencing with Quintara Biosciences or Genewiz. Base editing and indel efficiencies were quantified using the EditR and ICE (EditCo) online tools, respectively.

### Pooled TNF screen

Primary monocytes from two healthy donors were isolated as described, and 360x10^6^ cells from each donor were plated in 12-well plates (2.5x10^6^ cells in 2 ml of media per well) in SF macrophage media + 20 ng/ml M-CSF. Additional culture media was either added or changed every 48 hours. 3 days after isolation, monocyte-derived macrophages were transduced with a sgRNA library packaged in a Vpx-containing lentiviral vector with a GFP marker at a pre-defined dose (7.5 ul of concentrated virus in 1 ml of media per well), resulting in ∼65% and 90% transduction rate for either of the donors, respectively. 24 h after lentiviral transduction, the macrophages were treated with eVLP particles containing a scrambled/non-target sgRNA (50 ul of concentrated eVLP in 1 ml of media per well) and subsequently cultured in media with M-CSF as described above for another 5 days. Macrophages were stimulated with 50 ng/ml of ultrapure LPS isotype O:55 (Enzo) in the presence of GolgiPlug reagent (BD Biosciences) for 5 h. Cells were then detached with Accutase for 30 min at 37°C (Corning), washed once with EasySep buffer and stained with Ghostdye Violet450 (Tonbo) at 1:1000 dilution in EasySep buffer for 30 min on ice. Cells were then fixed using the CytoFix/Cytoperm kit (BD Biosciences; 200 ul of Cytofix solution per 1 million cells) and stained with a APC-labeled human TNF-alpha (Biolegend). Cells were washed once with Perm/Wash buffer and resuspended at 20x10^6^ cells/ml in EasySep buffer. Cell sorting was performed on a FACSARIA cell sorter at the UCSF Parnassus Flow Cytometry Core facility. We gated the population of single cells, which are GFP positive (transduced with lentivirus) and Ghostdye negative (viable) population and sorted the top and bottom 20% of TNF-alpha expressing cells. At least 12 million cells were sorted into each bin ensuring sufficient coverage.

### Pooled CD80 screen

Primary monocytes from two healthy donors were isolated as described, and 360x10^6^ cells from each donor were plated in 12-well plates (2.5x10^6^ cells in 2 ml of media per well) in SF macrophage media + 20 ng/ml M-CSF. Additional culture media was either added or changed every 48 hours. 4 days after isolation, monocyte-derived macrophages were transduced with the transcription factor and immunomodulator sgRNA library packaged in a Vpx-containing lentiviral vector with a GFP marker at a pre-defined dose (15 uL of 100x concentrated virus in 1 ml of media per well), resulting in 70% transduction rate for both donors. 24h after lentiviral transduction, the macrophages were treated with eVLP particles containing a scrambled/non-target sgRNA (35 ul of concentrated eVLP in 1 ml of media per well) and subsequently cultured in media with M-CSF as described above for another 5 days. Macrophages were stimulated with 50 ng/ml of ultrapure LPS isotype O:55 (Enzo) in the presence of GolgiPlug reagent (BD Biosciences) for 5 h. Cells were then detached with Accutase for 30 min at 37°C (Corning), washed once with EasySep buffer and stained with Ghostdye Violet450 (Tonbo;1:1000) and APC anti-CD80 (Biolegend; 2 ul of antibody per 1 million cells) in EasySep buffer for 30 min on ice. Cells were washed once with Easysep buffer and resuspended at 20x10^6^ cells/ml in EasySep buffer. Cell sorting was performed on a FACSARIA cell sorter at the UCSF Parnassus Flow Cytometry Core facility. We gated on single cells, which are GFP positive (transduced with lentivirus) and Ghostdye negative (viable) and sorted the top and bottom 20% of CD80 expressing cells. At least 10 million cells were sorted into each bin ensuring sufficient coverage.

### Genomic DNA extraction

Sorted cells were pelleted and resuspended at up to 5x10^6^ cells per 400 µl of lysis buffer (1% SDS, 50 mM Tris, pH 8, 10 mM EDTA). The remaining protocol reflects additives/procedures performed per each 400 µl reaction. 16 µl of NaCl (5M) was added, and the sample was incubated on a heat block overnight at 66°C. The next morning, 8 µl of RNAse A (10mg/ml; Thermo Fisher; FEREN0531 was added, and the sample was vortexed briefly, and incubated at 37°C for 1 hour. Next, after 8 µl of Proteinase K (20mg/ml) (ThermoFisher; AM2548) was added, the sample was vortexed briefly and incubated at 55°C for 1 hour. Homemade phase lock tubes were prepared for each sample by spinning 200 µl sterilized high vacuum grease gel to the bottom of a 1.5 ml tube at 20,000g for 1 minute and then 400 µl of Phenol:Chloroform:Isoamyl Alcohol (25:24:1) was added to each tube. 400 µl of the sample was then added to the phase lock tube and the tube was shaken vigorously. The sample was centrifuged at maximum speed at room temperature for 5 minutes. The aqueous phase was transferred to a low-binding eppendorf tube (Eppendorf, Cat #022431021) and then 40 µl of Sodium Acetate (3M), 1µl GlycoBlue (Invitrogen, Cat # AM9515), and 600µl of room temperature isopropanol was added. The sample was then vortexed and stored at -80°C for 30 minutes or until the sample had frozen solid. Next the sample was centrifuged at maximum speed at 4°C for 30 minutes, the pellet was washed with fresh 70% room temperature Ethanol, and allowed to air dry for 15 minutes. Pellets were then resuspended in Zymo DNA elution buffer (Zymo, Cat No: D3004-4-10), and placed on the heat block at 65°C for 1 hour to completely dissolve the genomic DNA.

### Library preparation for next generation sequencing

sgRNA cassette was amplified and barcoded from the genomic DNA as initially described by Joung et al^71^. Up to 2.5 µg of genomic DNA were added to each 50 µL reaction, which included 25 µL of NEBNext Ultra II Q5 master mix (NEB, Cat #M0544L), 1.25 µL of the 10 µM forward primer and 1.25 µL of the 10 µM reverse primer, and H2O to 50 uL. Due to differences in sgRNA scaffold in the respective plasmid backbone, different reverse primer sets were used for CD80 and TNF screens. The following PCR cycling conditions were used: 98°C for 3 minutes, followed by 23 cycles at 98°C for 10 seconds, 63°C for 10 seconds, and 72°C for 25 seconds, and ending with 2 minutes at 72°C. Amplicons were purified using SPRIselect beads following manufacturer’s protocols (Beckman). The concentration of each sample was then measured using the Qubit dsDNA high sensitivity assay kit (Thermo Fisher Scientific, Cat #Q32854). Samples were mixed at equimolar ratios, diluted to 750 pM in RSB + Tween (Illumina) and PhiX (Illumina, #15017872) was spiked in to ∼10-20%. Libraries were sequenced on an Illumina NextSeq2000 instrument equipped with a P1 (100 cycles) X-LEAP-SBS cartridge (Illumina, Cat #15017872) with a custom sequencing primer diluted to 0.3 µM in HP21 (Illumina). Primer sequences are listed below (N indicates position of unique barcodes):

P5 primer (used for both TNF and CD80 screen):

AATGATACGGCGACCACCGAGATCTACACGCTTTATATATCTTGTGGAAAGGACGAAACAC C

P7 Primer(s) used for TNF screen:

CAAGCAGAAGACGGCATACGAGATNNNNNNNGTGACTGGAGTTCAGACGTGTGCTCTTCC GATCT CTAATAGGTGAGCggccAAGTTGATAACG

P7 Primer(s) used for CD80 screen: CAAGCAGAAGACGGCATACGAGATNNNNNNNNGTGACTGGAGTTCAGACGTGTGCTCTTCCGATCTctttgctgtttccagcaaagttgataacg Custom sequencing primer:

CCGAGATCTACACGCTTTATATATCTTGTGGAAAGGACGAAACACC

### Pooled screen analysis

A table of individual guide abundance in each sample was generated using the count command in MAGeCK v.0.5.9.5^72^. The MAGeCK test command was used to identify differentially enriched sgRNA targets between the low and high bins. All genes with an FDR-adjusted *P* < 0.05 were considered significant.

### Immunoblotting

Cells were seeded, differentiated and stimulated as indicated in a 24-well plate. Media was aspirated from cells and cells were lysed by adding 200 μl of 1x Laemmli buffer (BioRad) + 1% Beta-mercaptoethanol directly to the well. Lysates were transferred into tubes, DNA was sheared by passing solution through a 26-gauge needle using a syringe and samples were incubated at 65°C for 10 mins to facilitate denaturation of proteins. 25 μl of each sample were loaded on a NuPAGE Bis-Tris 4 - 12% gradient gels (ThermoFisher) and separated by SDS-PAGE in 1x NuPAGE MOPS running buffer at a constant voltage of 150V for 1 h. Proteins were transferred onto a PVDF membrane using Trans-Blot Turbo Mini PVDF Transfer packs according to manufacturer’s protocol (BioRad). Membranes were blocked with 5% milk in Tris-buffered saline with 0.05% Tween 20 (TBST) for 1h at RT. After washing three times with TBST for 5 min at RT, primary antibody solution was added and incubated overnight at 4°C. After washing three times with TBST for 5 min at RT, species-matched HRP-conjugated secondary antibody (Jackson ImmunoResearch) diluted in TBST + 5% milk was added to membranes followed by incubation at RT for 1h. After washing three times with TBST for 5 min at RT, ECL substrate (SuperSignal West Femto or Pico PLUS; Thermo Fisher) was added to membranes and chemiluminescence was visualized on a ChemiDoc imager (BioRad). The following primary antibodies were used in TBST + 5% BSA at a 1:1000 dilution: rabbit anti-CD80 (Cell Signaling; 15416S), rabbit anti-PD-L1 (Cell signaling; 13684T), rabbit anti-TNFAIP3 (Cell Signaling; 5630T), rat anti-actin (Biolegend; 664802), mouse anti-actin (MilliporeSigma; A5441).

### qRT-PCR

Cells were stimulated as indicated, lifted using accutase for 30 min at 37°C and collected into nuclease-free microcentrifuge tubes by centrifugation at 500 g for 5 min. Cell pellets corresponding to ∼0.5x10^6^ cells were snap-frozen in dry-ice and stored at -80°C until RNA extraction. RNA was extracted using the QuickRNA Micro-Prep kit (Zymo) according to the manufacturer’s protocols, RNA concentration was measured using a Nanodrop device and stored at -80°C. qRT-PCR was performed using the Taqman RNA-to-CT kit following manufacturer’s instructions (ThermoFisher) in 10 ul reactions containing 20 ng of input RNA in a 384-well format. PCR was run on a QuantStudio Real Time PCR system (ThermoFisher) under the following conditions: Step 1 – 48°C, 15 min, Step 2 – 95°C, 10 min, Step 3 – 95°C, 10 sec, Step 4 – 60°C, 30 sec, Step 5 – Go to step 3, 59X. CD80 RNA abundance was quantified using the ΔΔCT-method and normalized to the housekeeping gene GAPDH. The following gene expression assays were used: CD80 (Hs01045161_m1), GAPDH (Hs99999905_m1).

### AAV production

AAV6 cargo vector encoding HDR template sequence for GFP-CLTA knock-in was described before^40^. HDR template sequence for insertion of GFP at the N-terminus of Rab11a were designed as described before^37^, ordered as gBlock (Integrated DNA Technologies) with NotI restriction inserted at both sides and cloned into the AAV6 cargo vector by restriction enzyme digestion and ligation. AAV6 was produced as described before (Nyberg et al 2023). Briefly, 90% confluent HEK293 cells seeded in 150 mm dishes were transfected with 6 μg of cargo vector, 8 μg of Rep-Cap plasmid, and 11 μg of adenovirus helper plasmid in 200 μl of PEI. Approximately 72 hours post-transfection, cells were harvested in AAV lysis buffer (50 mM Tris, 150 mM NaCl), lysed via three rapid freeze-thaw cycles, and incubated at 37°C for 1 h with Benzonase (25 units/ml, Millipore Sigma #70-664-3). After cell harvest and PEG precipitation of the supernatant, AAV vectors were purified using iodixanol (OptiPrep, STEMCELL Technologies #07820) gradient ultracentrifugation. Vector titers were quantified by qPCR following DNase I (NEB #B0303S) treatment and Proteinase K (Qiagen #1114886) digestion, using primers specific to the viral genome. qPCR reactions were conducted with SsoFast EvaGreen Supermix (Bio-Rad #1725201) on a StepOnePlus Real-Time PCR System (Applied Biosystems #4376600). Titers were calculated by comparison with a serially diluted plasmid DNA standard of known concentration.

### Site-specific Knock-in

For site-specific knock-in, cells were co-treated with 20-50 µL of Cas9EDV and AAV6 encoding target-matched HDR template at a multiplicity of infection of 5e5. For site-specific knock-in in primary human macrophages, 0.5x10^6^ monocytes per well were plated in 24-well plates and co-treated with EDV and AAVs on Day 4 post-isolation in 500 µL of SF macrophage media supplemented with M-CSF or GM-CSF. Spinfection was performed at 1250 g and 32°C for 1 h. Media was changed to fresh SF macrophage media + cytokines after 24h, and cells were cultured for 9 additional days before analysis by flow cytometry. THP-1 and U937 cells were cultured in RPMI supplemented with 10% FCS, 1x L-Glutamine, 1x non-essential amino acids, Penicillin/Streptomycine, and sodium pyruvate (hereafter referred to as complete RPMI) and maintained at a density between 0.1x10^6^ and 1x10^6^ cells/ml for routine maintenance. 0.5x10^6^ cells were co-treated with 50 µl of Cas9EDV and AAV6 encoding target-matched HDR template at a multiplicity of infection of 500,000 in 500 µl of complete RPMI with polybrene (1:2000; EMD Millipore) in a 24-well plate. Spinfection was performed at 1250 g and 32°C for 1 h. 24 h post-infection, media was swapped to fresh cRPMI and cells were cultured for at least 5 days before checking GFP expression by flow cytometry. To establish cell lines with homogeneous GFP expression, top 5-10% highest GFP-expressing cells were sorted on a BD FACSAria II cell sorter.

### sgRNA library cloning

For the library used in the TNF screen, candidate genes were selected based on a previously published study^53^. Corresponding sgRNA sequences were obtained from the Brunello library^73^. Custom oligonucleotide pools, incorporating additional nucleotides containing the BsmBI-v2 recognition site, were synthesized (Twist Bioscience). Oligo pools were amplified using 10 cycles of PCR with KAPA HiFi polymerase following the manufacturer’s instructions (Twist). PCR amplicons were purified with SPRI beads, and the size was confirmed via E-gel electrophoresis. Golden Gate assembly was performed using 500 fmol of PCR inserts and 500 fmol of BsmBI-v2–digested LRG2.1-GFP backbone with an improved sgRNA scaffold described previously^59^. The assembled library plasmids were electroporated into *E. coli* MegaX DH10B T1R electrocompetent cells (C640003, Thermo Fisher) using a Bio-Rad Gene Pulser Xcell Electroporator with the following parameters: 2.0 kV, 200 Ω, and 25 μF. Immediately after pulsing, 1 mL of recovery medium was added and transferred into 2 mL of additional recovery medium, followed by incubation at 37 °C for 1 h. Cultures were then transferred into 200 mL LB medium for large-scale plasmid preparation according to the manufacturer’s protocol (Qiagen Maxiprep kit). Following confirmation that the desired colony-forming units (c.f.u.) per construct were achieved, 50 ng of library plasmid was subjected to PCR for indexing as described under ‘Library preparation for next generation sequencing’. PCR products were quantified using the Qubit dsDNA High Sensitivity Assay Kit (Thermo Fisher) and sequenced on a NextSeq P2000 platform (Illumina). sgRNA abundance was quantified using the count command in MAGeCK v 0.5.9.5. Composition and cloning of the transcription factor and immunomodulator library used in the CD80 screen was described previously^59^. The TF and immunomodulator library was in a LRG2.1-GFP backbone without scaffold modifications.

### CAR-macrophage killing assay

CD14⁺ monocytes were isolated from Leukopaks using the monocyte isolation enrichment kit (StemCell Technologies) as described above. Monocytes were cultured in ImmunoCult SF Macrophage Medium (StemCell Technologies) supplemented with 10 ng/mL GM-CSF(Peprotech) for 4 days. On day 4, cells were transduced with LentiVPX Her2CAR and eVLPs, followed by a medium change to cRPMI (10% FBS, 1% penicillin–streptomycin, 10mM HEPEs) containing 10 ng/mL GM-CSF and spinfection (1,250g, 32°C, 30 min). One day after transduction, the medium was completely replaced, and cells were maintained for an additional 3 days. SK-OV3 cells stably expressing mKate were plated one day before co-culture with CAR-macrophages in cRPMI without GM-CSF, and co-cultures were monitored using an Incucyte imaging system

### Cytokine measurement

To quantify inflammatory cytokines, we used either the LEGENDplex Human Inflammation Panel 1 (BioLegend) or the Human Inflammatory Cytokines Cytometric Bead Array Kit (BD Biosciences) according to the manufacturers’ protocols. Primary human macrophages, either nucleofected or treated with virus-like particles as described above, were fully differentiated into macrophages for 7 days. Subsequently, macrophages were detached, re-plated in 96-well plates at equal cell densities across experimental groups, and allowed to recover for 24 h. Cells were then stimulated with 100 ng/mL LPS and 20 ng/mL IFN-γ for an additional 24h. Culture supernatants were collected, clarified by brief centrifugation, and stored at −80 °C until analysis.

## Acknowledgements

We thank all members of the Eyquem, Marson and Carnevale laboratory for their support, input, and technical assistance. We also thank V. Nguyen for technical assistance with flow cytometry and cell sorting at UCSF Parnassus Flow core; S. Dodgson and N. Leung for scientific advice and writing; T. Tolpa for graphical polishing; C. Ward for sgRNA design for base editing. N.L. was funded by the NIH (grants DA046100 and AI122390). J.K.N. and D.X. acknowledge funding for this work from the NIH (R35GM155044), Laboratory for Genomics Research, and the Bakar Fellows Program. D.X. is funded by the Weill Neurohub Fellows Program. M. O. is supported by the Astellas Foundation for Research on Metabolic Disorder and the Chugai Foundation for Innovative Drug Discovery Science. J.C. was supported by NIH/National Cancer Institute K08, 1K08CA252605-01, a Burroughs Wellcome Fund Career Award for Medical Scientists, the Lydia Preisler Shorenstein Donor Advised Fund, the Parker Institute for Cancer Immunotherapy, and the Pascarella Scholars Fund. A.M. received funding from the Simons Foundation, Lloyd J. Old STAR Award (Cancer Research Institute), Parker Institute for Cancer Immunotherapy, Innovative Genomics Institute, Larry L. Hillblom Foundation (grant 2020-D-002-NET), Northern California JDRF Center of Excellence, the Byers family, K. Jordan and the CRISPR Cures for Cancer Initiative. P.D. is a Fellow of The Jane Coffin Childs Fund for Medical Research. H.J. is a Parker Institute for Cancer Immunotherapy Scholar Awardee and was supported by the National Research Foundation of Korea (NRF) through the Postdoctoral Overseas Training Program (RS-2023-00242661).

## Author information

### Contributions

H.J., P.D., A.M., and J.C. conceptualized the study. H.J. and P.D. led and performed all experiments with C.C. and E.U. providing support. H.J., M.O., and Z.S. performed Bulk-RNA sequencing and computational analysis. L.S. and J.H.J. produced AAV6. J.H., W.N., J.A.D., D.X., J.K.N., V.A., T.T., N.L., M.A., D.R.L., and J.E. provided and characterized resources that were used in this study. H.J., P.D., A.M., and J.C. wrote the manuscript with input from all authors

---

**Extended Data Figure 1.**
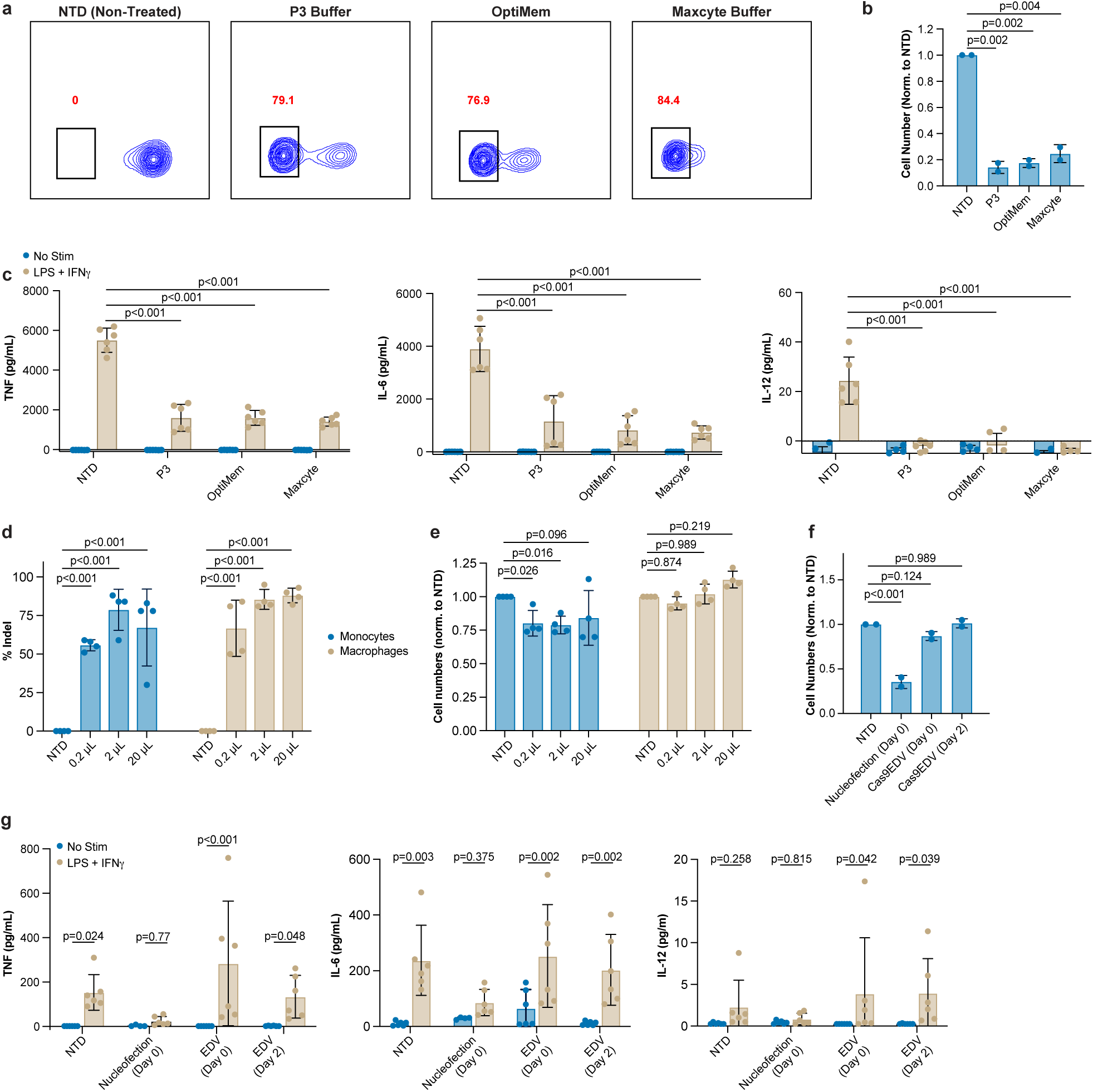
: VLP-mediated gene editing, but not nucleofection maintains viability of human primary myeloid cells. **a**, Representative flow cytometry plots showing knockout of B2M protein in primary human macrophages after delivery of Cas9-RNP by nucleofection in indicated buffers. Nucleofection was performed Day 0 post-isolation and cells were analyzed by flow cytometry 7 days later. **b**, Yield of macrophages that underwent nucleofection in indicated buffers, normalized to the non-treated condition. **c**, Nucleofected primary human macrophages were stimulated with 100 ng/ml of LPS and 25 ng/ml of IFN-γ and cytokine release was measured after 24 h. **d**,**e**, Quantification of macrophages or monocytes with indels at the sgRNA target site after B2M Cas9-eVLP treatment of monocytes or macrophages. (**d**) and yield of macrophages treated with Cas9eVLP, normalized to the non-treated condition (**e**). **f**, Yield of dendritic cells that underwent nucleofection in indicated buffers on Day 0, normalized to non-treated condition. **g**, Nucleofected primary human dendritic cells were stimulated with 100 ng/ml of LPS and 25 ng/ml of IFN-γ and cytokine release was measured after 24 h. One-way ANOVA followed by means of Dunnett’s multiple comparison (**b**,**d,f**); two-way ANOVA followed by means of Tukey’s multiple comparison (**c**,**g**)

**Extended Data Figure 2.**
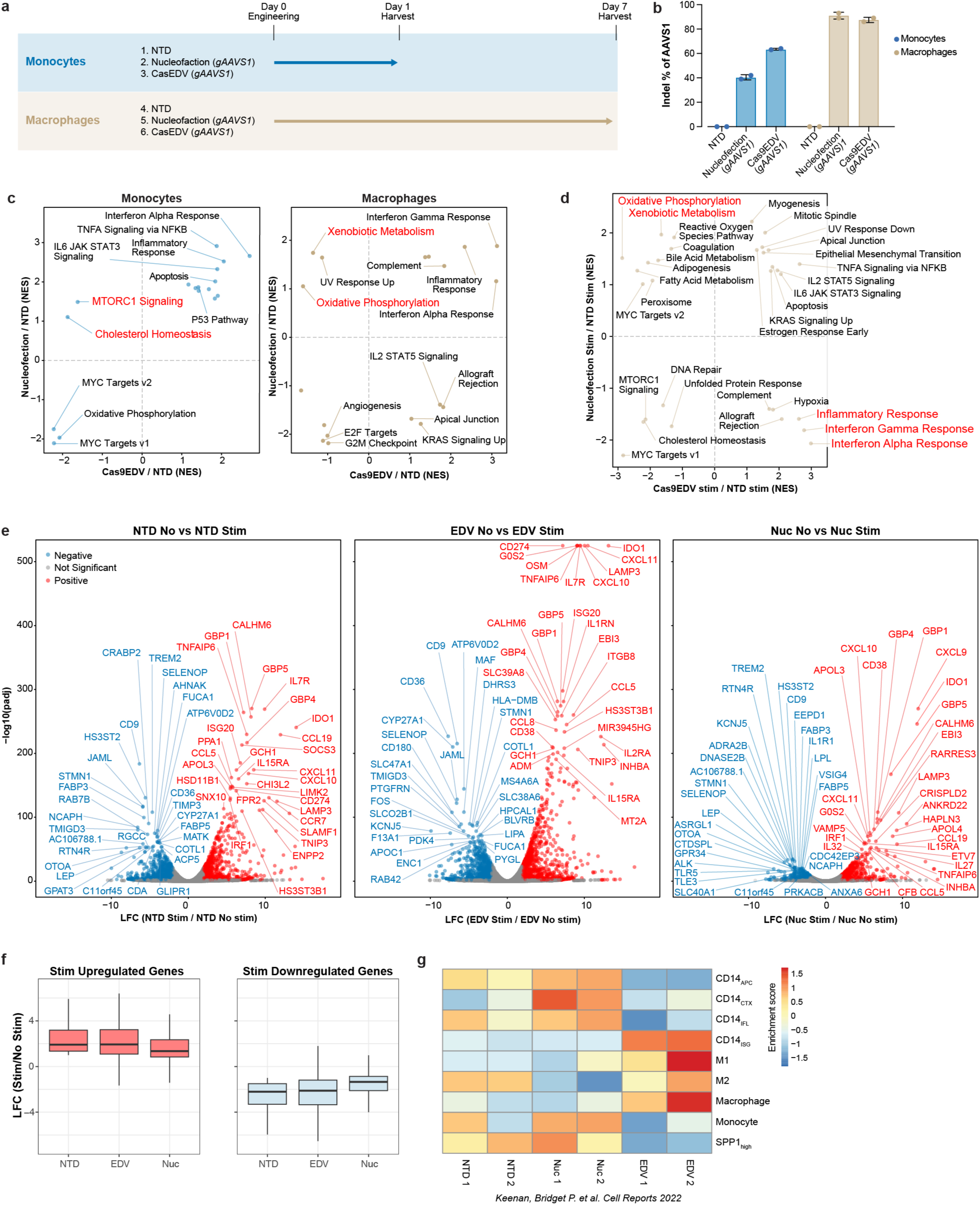
Bulk RNA-seq analysis uncovers transcriptional changes following nucleofection and Cas9EDV treatment. **a**, Workflow for Bulk RNAseq experiments. Human primary monocytes or macrophages were gene treated with AAVS1-targeted Cas9EDVs or left non-treated before isolation of total RNA and Bulk transcriptomics analyses. **b**, Analysis of indels at AAVS1 locus in macrophage cells used in Bulk RNAseq experiments. Genetic locus around sgRNA target site was amplified by PCR and analyzed by ICE analysis. **b**, Gene Set Enrichment Analysis (GSEA) of bulk RNA-seq from primary monocytes and macrophages, comparing nucleofection or Cas9EDV treatment to non-treated (NTD) controls. Significantly enriched pathways are shown on plots. **d**, Heatmap of Gene Set Variation Analysis (GSVA) scores projected onto previously reported, established monocyte/macrophage signatures: CD14APC (antigen processing and presentation), CD14CTX (chemotaxis molecules and suppressive cytokines), CD14IFL (inflammatory pathways), and CD14ISG (interferon-stimulated gene response). We compared the upregulated gene sets in nucleofected or Cas9EDV-treated macrophages with previously published transcriptional signatures of specific macrophage subpopulations. **e** Volcano plots showing differential gene expression with fold change and adjusted P value (Padj) for NTD, EDV-treated, and nucleofected samples upon LPS and IFN-γ stimulation, compared to their corresponding non-stimulated controls. Red indicates significant upregulation; blue indicates significant downregulation. NTD, non-treated; EDV, Cas9EDV-treated; Nuc, nucleofection. **f,** Bar plots showing the distribution in LFCs for defined upregulated or downregulated genes upon stimulation (∼1200 genes each). **g**, GSEA identifying pathways overlapping between Cas9EDV-treated and nucleofected groups, each normalized to the NTD group after 24 h LPS and IFN-γ stimulation. Significantly enriched pathways are shown on plots. Upregulation of genes involved in xenobiotic metabolism pathways observed in nucleofected cells after LPS/IFN-γ stimulation.

**Extended Data Figure 3.**
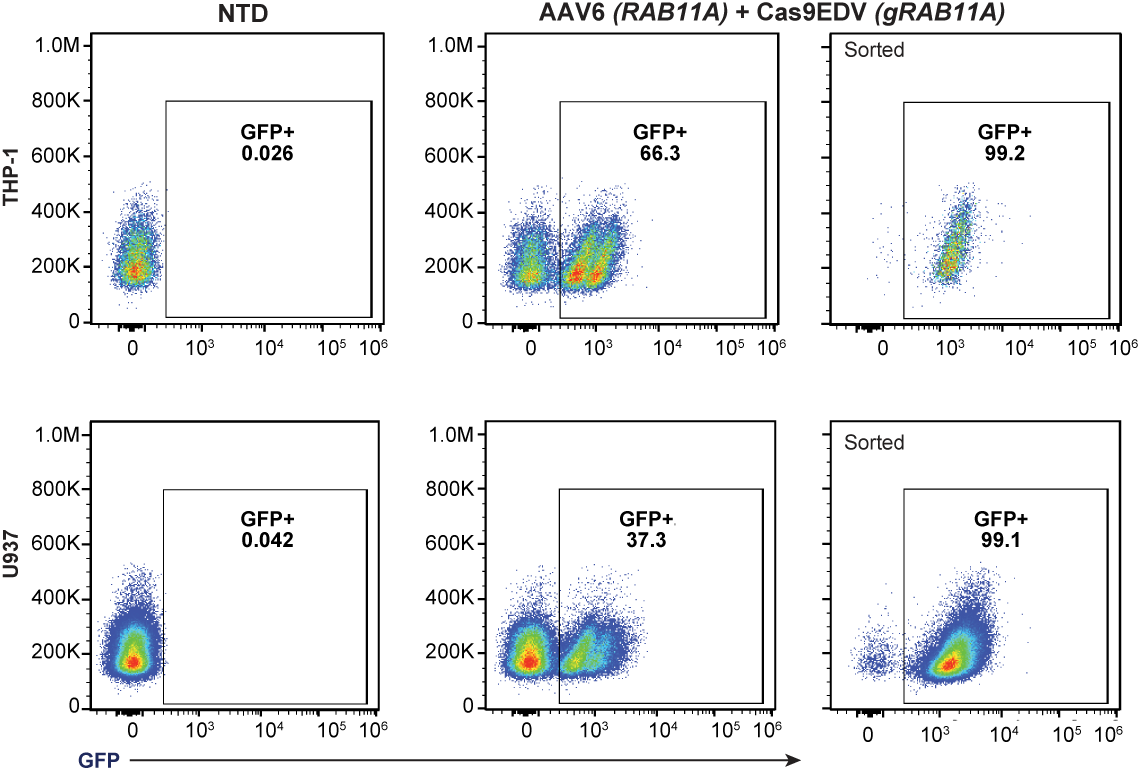
Cas9EDV/AAV co-treatment for site-specific integration in human myeloid cell lines. **a**, Flow cytometry analysis of THP-1 and U936 cells treated with Rab11a-targeted Cas9EDV and AAV6 encoding a GFP donor template. Successful knock-in was quantified by GFP fluorescence. GFP⁺ cells were sorted to establish stable cell lines with homogeneous GFP-Rab11a expression.

**Extended Data Figure 4.**
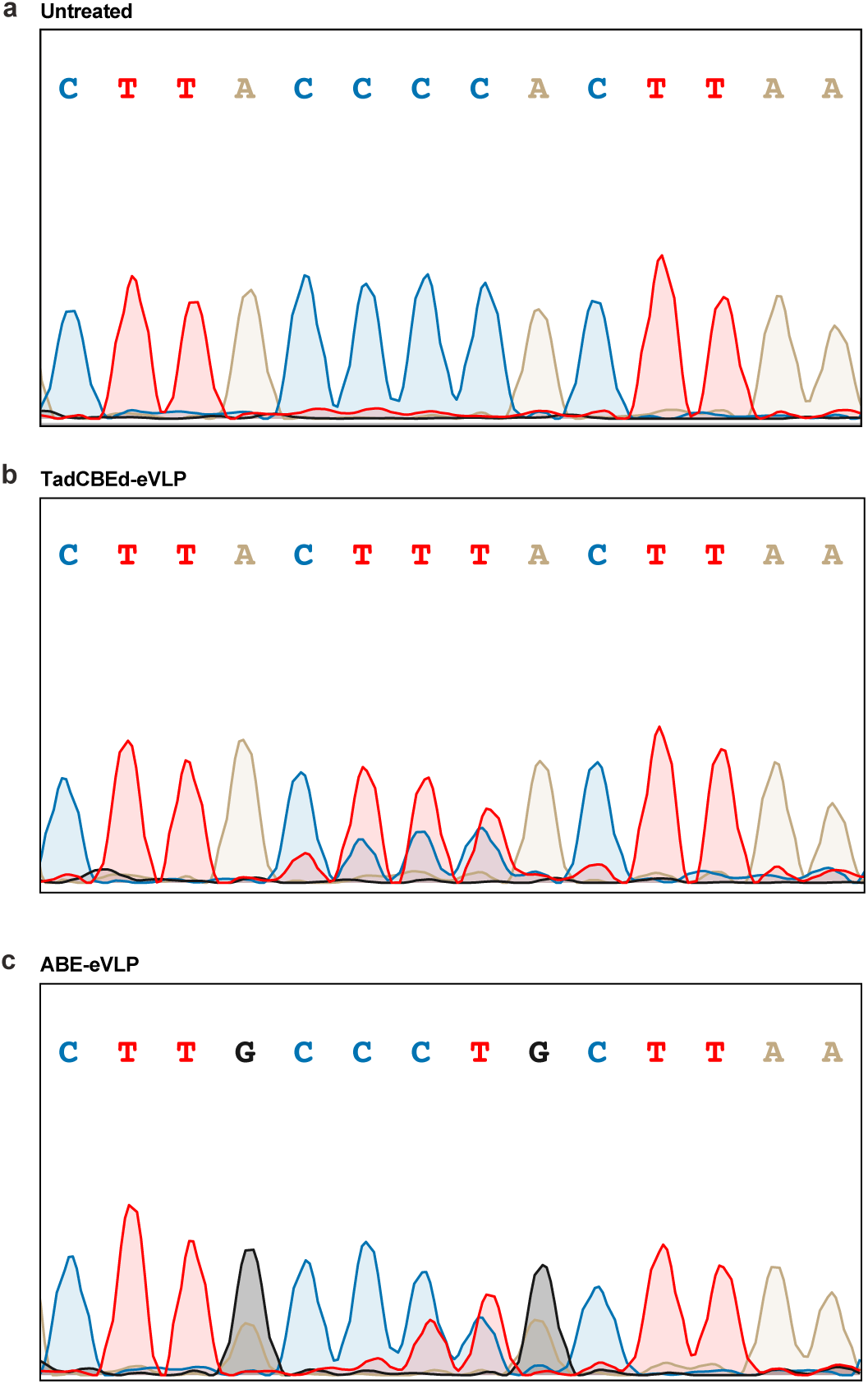
Analysis of eVLP-mediated base editing. **a**-**c**, Representative sanger sequencing traces of macrophages treated with ABE8e-eVLPs (**b**) or TadCBEd-eVLPs (**c**) loaded with guide targeting splice site in *B2M* gene.

**Extended Data Figure 5.**
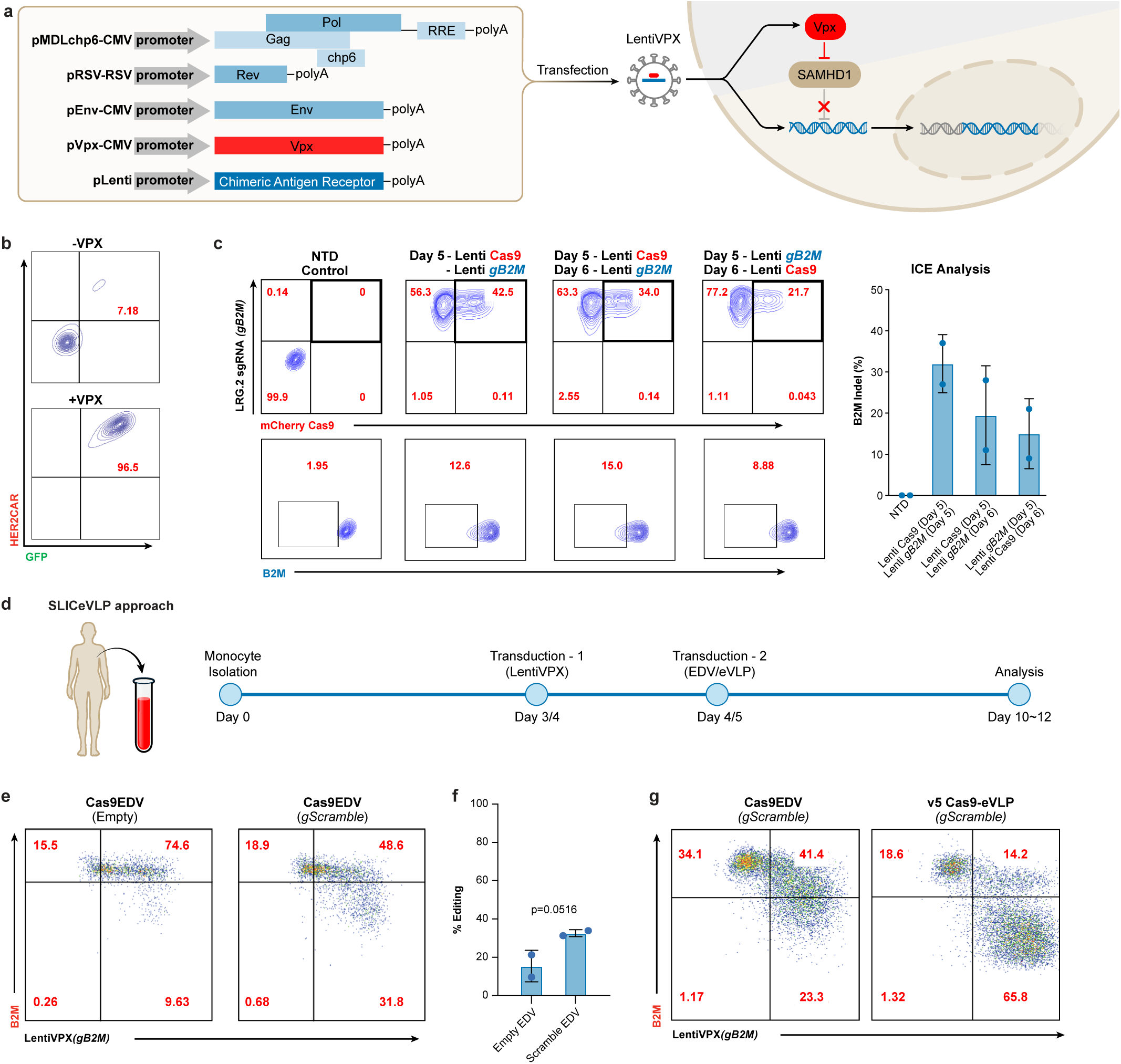
Optimization of SLICeVLP workflow. **a**, Schematic representation showing production of VPX-containing lentiviral vectors. **b**, Representative flow plots showing transduction efficiency of macrophages with HER2CAR lentiviral vector with and without VPX. **c**, Human macrophages were sequentially transduced with LentiVPX vectors encoding a *B2M-*targeting sgRNA (together with a BFP marker) and Cas9 (together with an mCherry marker). Representative flow plots and quantifications of indel generation at the genomic level are shown. **d**, Schematic representation of workflow for SLICeVLP optimization experiments. **e**, Representative flow cytometry analysis of macrophages transduced with VPX lentiviral vector encoding an sgRNA targeting B2M followed by treatment with empty or scramble sgRNA-loaded Cas9-EDVs. **f**, Quantification of B2M depletion. *P* values calculated by one-sided unpaired Student’s *t*-test. **g**, Representative flow cytometry analysis of macrophages transduced with Vpx lentiviral vector encoding an sgRNA targeting B2M followed by treatment with scramble sgRNA-loaded Cas9-EDVs or Cas9-eVLPs.

**Extended Data Figure 6.**
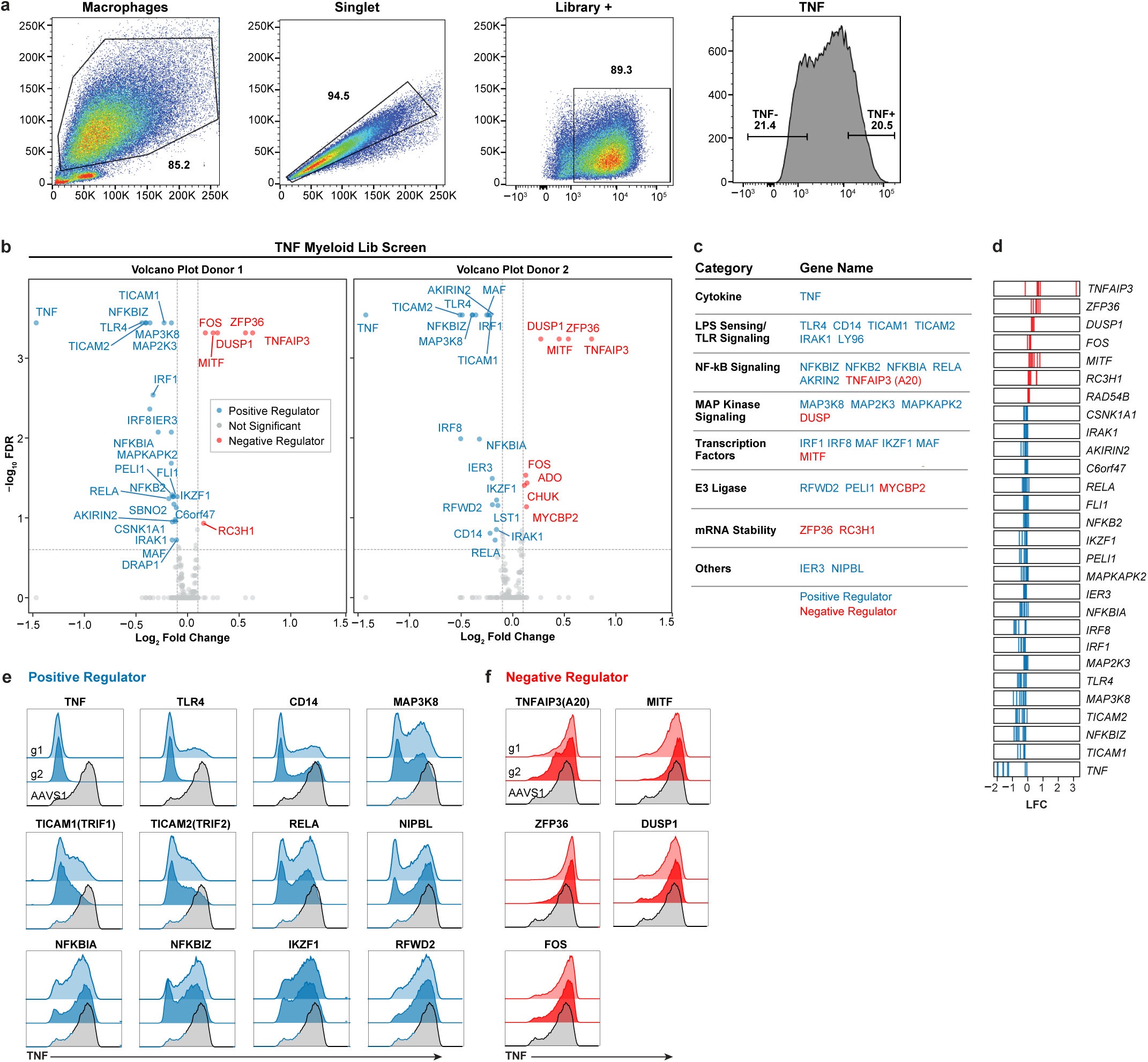
A pooled screen to uncover regulators of TNF production **a**, Flowcytometry gating strategies for TNF screen. **b**, Volcano plots for hits from TNF screen. X-axis shows mean LFC of all sgRNAs for each gene. Y axis shows -log false discovery rate (FDR). All hits with |LFC| > 0.1 are highlighted in red (negative regulators) or blue (positive regulators). Volcano plots from both biological replicates are displayed separately. **c**, Classification of selected hits from TNF screen based on known functions. **d**, Rug plots showing log fold change (LFC) for individual sgRNAs in TNF screen. **e**,**f**, Representative flow cytometry plots from arrayed validation experiments for TNF screen. Macrophages were treated with Cas9-eVLPs targeting the indicated positive (**e**) or negative (**f**) regulators, stimulated with 50 ng/ml LPS for 5 h before assessment of TNF production by intracellular cytokine staining and flow cytometry. Flow cytometry plots are representative of at least 4 independent experiments.

**Extended Data Figure 7.**
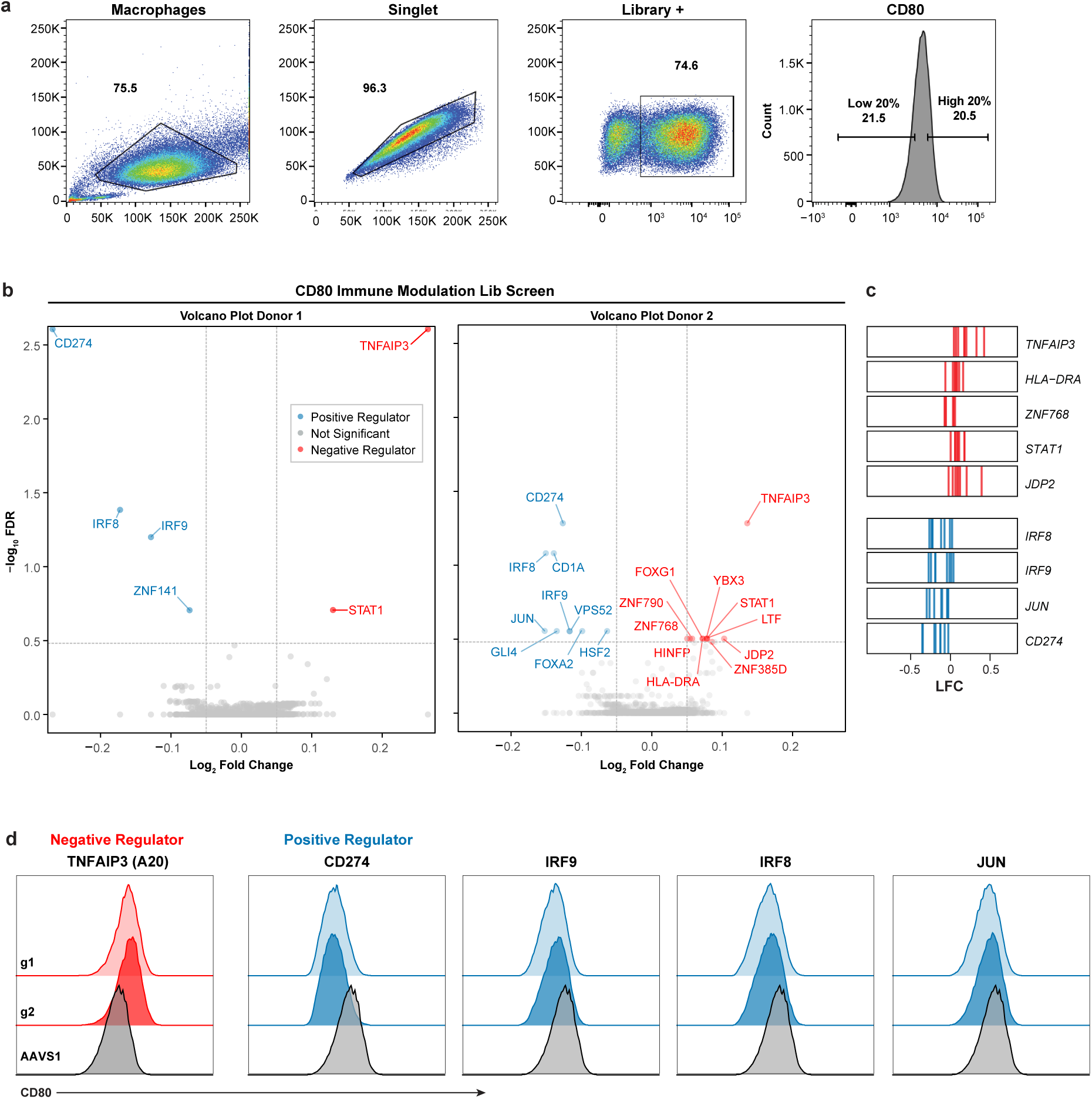
A pooled screen to uncover regulators of CD80 surface expression. **a**, Flowcytometry gating strategies for CD80 screen. **b**, Volcano plots for hits from CD80 screen. X-axis shows mean LFC of all sgRNAs for each gene. Y axis shows -log false discovery rate (FDR). All hits with |LFC| > 0.05 are highlighted in red (negative regulators) or blue (positive regulators). Volcano plots from both biological replicates are displayed separately. **c**, Rug plots showing log fold change (LFC) for individual sgRNAs in CD80 screen. **d**, Representative flow cytometry plots from arrayed validation experiments for CD80 screen. Macrophages were treated with Cas9-eVLPs targeting the indicated negative or positive regulators before assessment of CD80 surface levels by flow cytometry. All flow cytometry plots are representative of at least 4 independent experiments.

**Extended Data Figure 8.**
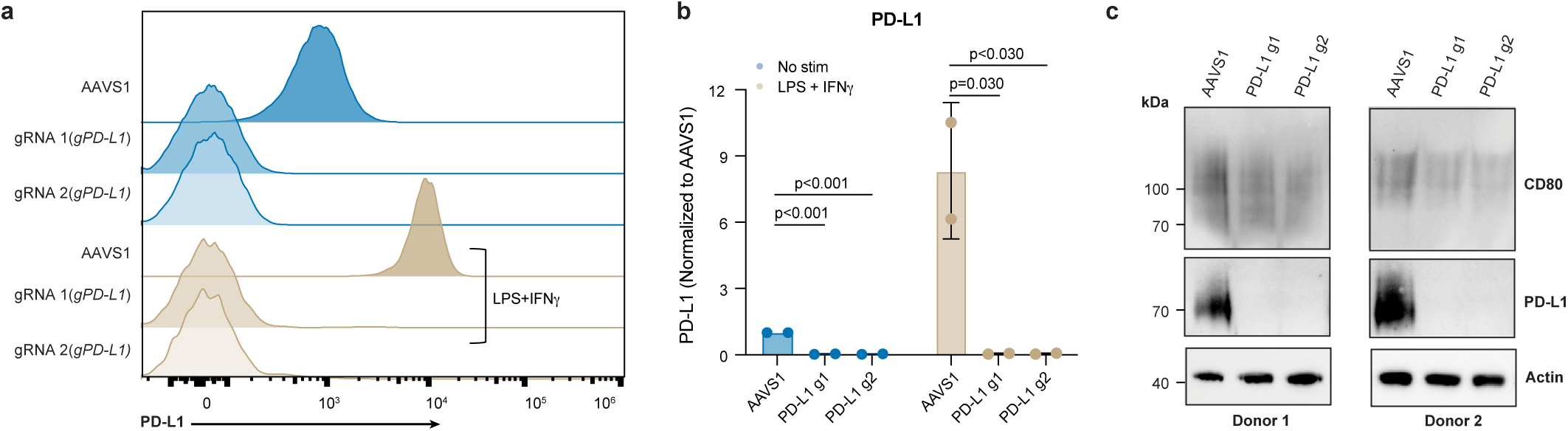
Validation of PD-L1 knockout in human primary macrophages and its effect on CD80 protein levels. **a**,**b**, Flow cytometry analysis of surface PD-L1 expression in macrophages treated with Cas9-eVLPs targeting CD274/PD-L1 under basal and LPS+IFN-γ stimulated conditions. Quantification of two independent experiments is shown in (**b**). *P* values calculated by means of Dunnett’s multiple comparison test after ordinary one-way ANOVA tests. **c**, Immunoblot showing levels of PD-L1 and CD80 in whole cell lysates from macrophages treated with Cas9-eVLPs targeting CD274/PD-L1 or AAVS1. Results from two human donors are shown.

**Extended Data Figure 9.**
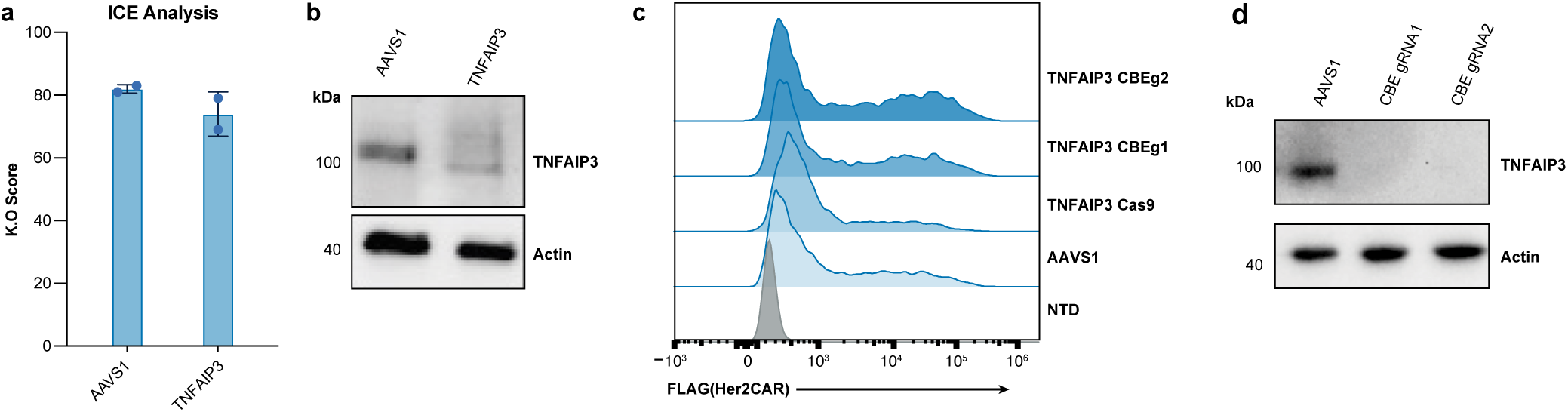
Characterization of TNFAIP3-ablated macrophages and HER2CAR macrophages. **a**, ICE analysis of genome editing efficiency at the AAVS1 and TNFAIP3 loci following Cas9EDV treatment. **b**, Immunoblot analysis of macrophages treated with Cas9-eVLPs carrying sgRNAs targeting *AAVS1* or *TNFAIP3*. **c**, Flow cytometry analysis of HER2CAR expression in primary human macrophages across experimental conditions where Cas9EDV or CBE-eVLP particles were used to disrupt *TNFAIP3.* **d**, Immunoblot analysis of macrophages treated with TadCBEd-eVLPs carrying sgRNAs designed to introduce a premature stop-codon in *TNFAIP3*.

